# Single-cell analysis of a salmonid immune system (brown trout *Salmo trutta*) reveals evolutionary divergence and hatchery-induced transcriptional reprogramming

**DOI:** 10.1101/2025.05.04.652114

**Authors:** James Ord, Helena Saura Martinez, Monica Hongroe Solbakken, Anastasiia Berezenko, Simone Oberhaensli, Stephanie Talker, Heike Schmidt-Posthaus, Irene Adrian-Kalchhauser

## Abstract

Vertebrate immune systems exhibit striking evolutionary diversity, yet our understanding remains biased toward mammalian models. Here, we generate a single-cell atlas of immune cells from the ecologically and economically important salmonid *Salmo trutta* (brown trout), a lineage characterized by an ancestral whole-genome duplication (WGD). Profiling over 83,000 kidney-derived immune cells, we resolved 34 transcriptionally distinct populations, identified core immune lineages, and uncovered novel markers in neutrophils, macrophages, T- and B-cells. We detected pervasive transcriptional divergence between WGD-derived ohnologue pairs, indicating putative sub- and neofunctionalization in immune gene regulation. We further show that the transcriptional identity of immune cells is shaped by rearing history: fish raised in hatcheries—whether for one or multiple generations—showed shifts in immune gene expression across cell types. These included genes involved in G protein-coupled receptor signalling, a process which has previously been implicated in domestication. Our findings provide insight into the evolution of vertebrate immunity and raise concerns about the immunological fitness of hatchery-reared fish released into the wild.

## Introduction

Vertebrate immune systems comprise some of the most complex systems in nature, exhibiting extraordinary diversity between species (Milgroom, 2023). Differences range from the structural level (e.g. in teleost fish, the kidneys are the principal source of immune cells, not bone marrow as in mammals), to the diversity of cell types (distinct cell types found in some species but not others), enzymes, and gene repertoire (Flies, 2020). New sequencing technologies, notably single-cell RNA sequencing now offer the opportunity to map the genetic regulation of vertebrate immune systems in unprecedented detail, including in non-model organisms. Such analyses can yield insights into evolutionary processes, from evolutionary arms races to the effects of gene duplications, and provide useful data for veterinary and human medicine. However, non-model organisms present unique analytical challenges, particularly those that are highly divergent from humans and model species (mostly mammals) from which most prior knowledge on gene function is derived. A case in point is the salmonid lineage, in which a whole genome duplication (WGD) event approx. 80MYA resulted in duplicated genes, many of which may have diverged in function thus making cell type identification difficult when relying on established markers from mammals.

Brown trout (*Salmo trutta*) are an ecologically and economically important member of the salmonid family of fishes (Lobón-Cerviá & Sanz, 2017). They are valued by recreational anglers and contribute significantly to the economically important sport fishing industry through tourism, sale of equipment, and access rights. They also play a crucial ecological role as apex predators in lotic alpine streams by controlling populations of smaller fish and insects and influence the overall structure and health of freshwater ecosystems through cascading effects. Based on their specific requirements regarding temperature, river structure, and water quality, they serve as sentinel species for riverine health. Brown trout populations suffer earlier than other species from pollution, habitat degradation, or changes in water temperature, and conservation efforts focusing on protecting brown trout habitats can have positive effects on the overall conservation of aquatic biodiversity. As a species of cultural and heritage value, brown trout also serve as flagship species to promote ecosystem conservation and protective management. At the same time, brown trout are under pressure from global change through the combined effects of changes in river structure, climate change, the translocation of competitor species and pathogens, and management practices. Their ability to mount appropriate immune responses is sensitive to temperature (Bailey et al., 2017; Strepparava et al., 2018) and is additionally impacted by water quality parameters (Rehberger et al., 2020; Schmidt-Posthaus et al., 2013). To support wild populations, large stocking programs are in place, yet the stocked fish fail to thrive (Baer et al., 2023), may be less resistant to pathogens, and the conditions experienced during raising offspring in artificial environments before release affect disease incidence (Palikova et al., 2017). This renders the immune system regulation of brown trout an important area of study.

The brown trout immune system is expected to largely follow the standard vertebrate build – i.e. innate and adaptive immune systems with many conserved cell types – but remains to be described in detail. Brown trout, as salmonids, have been subject to WGD followed by rediploidisation (return to disomic inheritance), accompanied by loss or retention of the resulting gene copies (Gillard et al., 2021; Macqueen & Johnston, 2014; Robertson et al., 2017). Amongst genes retained as duplicates, fates may include pseudogenization (because loss of function mutations are less detrimental), neofunctionalization (divergence of function), and sub-functionalization (division of roles in the same function) between the retained copies. For the related salmonid rainbow trout, only one quarter of 6733 pairs of WGD-derived gene duplicates (termed ‘ohnologs’) showed evidence of conserved function between the two genes as indicated by highly correlated expression patterns and similar expression levels across tissues (Berthelot et al., 2014).

Phylogenetically, brown trout is located at an interesting intermediate position between the well-studied whitefishes (Coregonidae) and trouts (Oncorhynchidae) (Crête-Lafrenière et al., 2012). They share a common ancestor with salmon (*Salmo Salar*) around 12 Mya, and with rainbow trout (Oncorhynchus mykiss) around 30 Mya (Crête-Lafrenière et al., 2012; Lecaudey et al., 2018). However, specific taxonomic relations, history of hybridisation and isolation, and even a potential multispecies taxonomy for brown trout populations are currently being discussed (Hashemzadeh Segherloo et al., 2021; Marić et al., 2023; Veličković et al., 2023). Understanding patterns of neofunctionalization in brown trout thus has the potential to complement existing data and provide perspective with regard to species-specific versus universal effects of WGD.

The immune system in fish features particularly interesting evolutionary trajectories. Teleost immune gene repertoires are highly specialized for each species, and feature prevalent non-canonical features compared to well-studied model systems such as loss of MHCII, novel subfamilies within NLR and TLR pattern recognition receptor families and loss of antibodies (Malmstrøm et al., 2016; Mirete-Bachiller et al., 2021; Solbakken et al., 2017; Suurväli et al., 2022; Swann et al., 2020). Therefore, the annotation of immune genes in fish can be challenging from genome sequence alone, where gene models may be incomplete or lack gene identification due to sequence divergence, or be misannotated due to highly variable gene families. For brown trout, there is a high-quality reference genome which would provide a solid foundation for downstream analyses (Hansen et al., 2021) but detailed functional annotations are lacking. Recent attempts to characterize the immune gene repertoire of brown trout have revealed extensive gene duplications in many pathways involved in immune cell differentiation and proliferation, suggesting many duplicates of immune-related genes have indeed been retained following WGD (Colgan et al., 2021). The functional implications of these duplications remain to be determined.

This study provides baseline data for the healthy brown trout cellular immune system using single cell RNA sequencing of enriched kidney-derived mononuclear cell populations from nine *S. trutta* individuals and analyses the impact of long-term evolutionary processes (whole genome duplication) and short-term environmental exposures (recent rearing history) on cellular immunity. We use single cell RNA sequencing of brown trout from three different sources to a) describe the immune cell types of brown trout, b) identify novel markers for particular cell types, c) investigate expression patterns of immune gene ohnologues to identify neofunctionalizations, and d) describe the effects of rearing environment on cellular immunity. We identify cell types based on previously established marker genes, explore the identity of cell types with non-canonical gene expression, and explore their gene expression patterns, and thus offer a set of marker genes for brown trout to use in future RNA-based analyses of immune functions in the absence of highly specific antibodies. To explore evolutionary peculiarities in brown trout, we analyse expression patterns of salmonid-specific ohnologues and propose functional diversification patterns for some of them. To investigate the impact of short-term environmental challenges on immune function, we compare three groups of brown trout with natural vs farmed rearing histories in terms of gene expression profiles at the cluster-level.

## Results

### Identification of cell lineages and subtypes

Single cell RNA sequencing from 83,847 hematopoietic cells, enriched from nine independent brown trout kidneys (**Fig. 1A**), identified 34 putative immune cell types. An initial set of 29 clusters (**Fig. S1**) was obtained from the top 2000 variably expressed genes and could be further refined to 34 final clusters based on the expression of lineage markers (**Fig. 1B**). 31 of these clusters could be assigned to the major hematopoietic cell lineages based on expression modules of ‘prior markers’ (**Fig. 2A**; **Table 1**: variably expressed prior markers; **Table S1**: full list of prior markers). The vast majority of cells assigned as putative neutrophils and B-cells, respectively (**Fig. 2B**).

**Figure 1.**
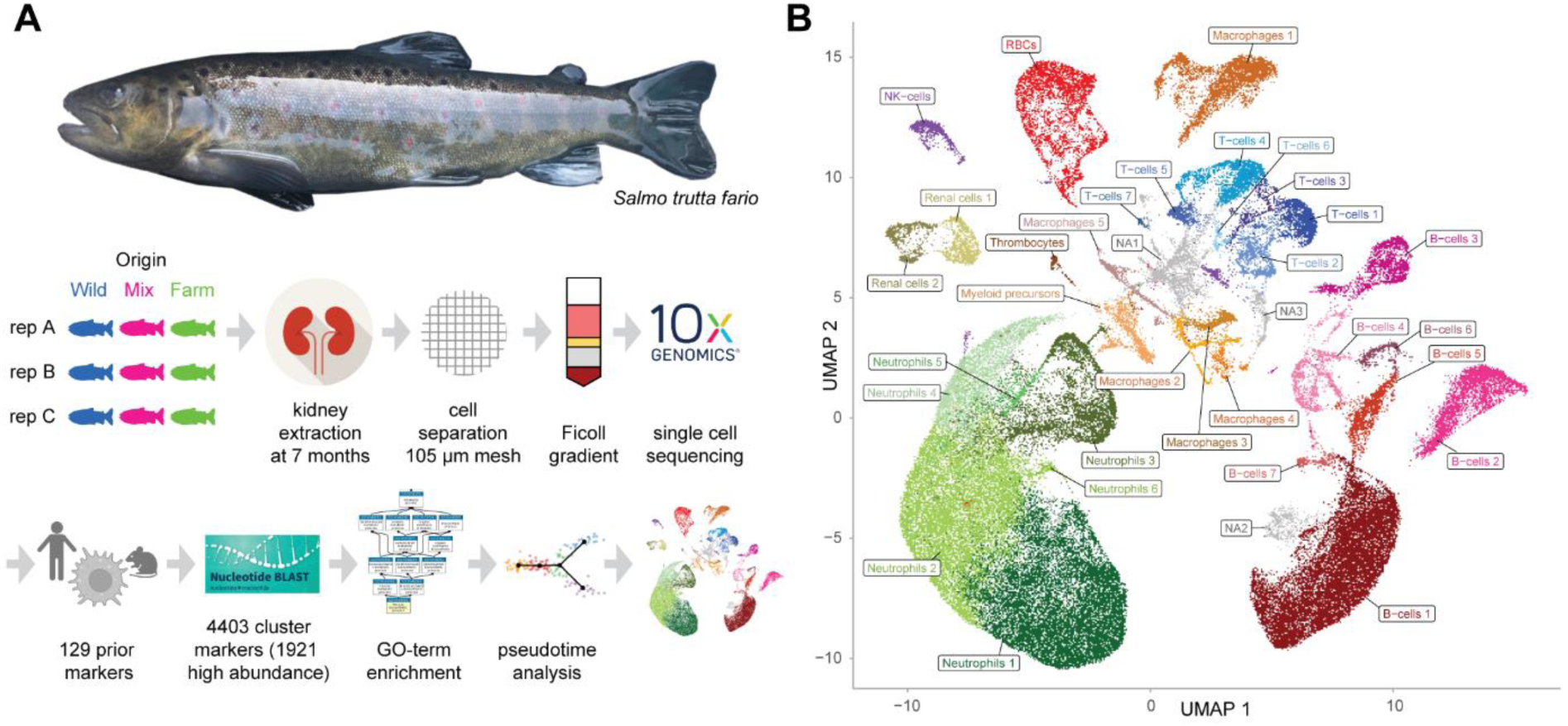
The brown trout cellular immune system. (A) Analysis setup. Kidney tissue was dissected from 9 juvenile Salmo trutta of 7 months of age, three each raised in the wild, in mixed conditions, and at a farm. Hematopoietic cells were extracted and used for single-cell RNAseq (scRNAseq). Cluster identity was assigned based on prior markes, newly identified cluster markers, GO terms, and pseudotime analysis. **(B) 34 immune cell clusters.** UMAP projection of 83,847 cells grouped into 34 clusters according to shared gene expression variation. Of the 34 clusters, 31 were assigned putative identities, while 3 remain unassigned. NA2 may represent phagocytic B-cells.

**Figure 2.**
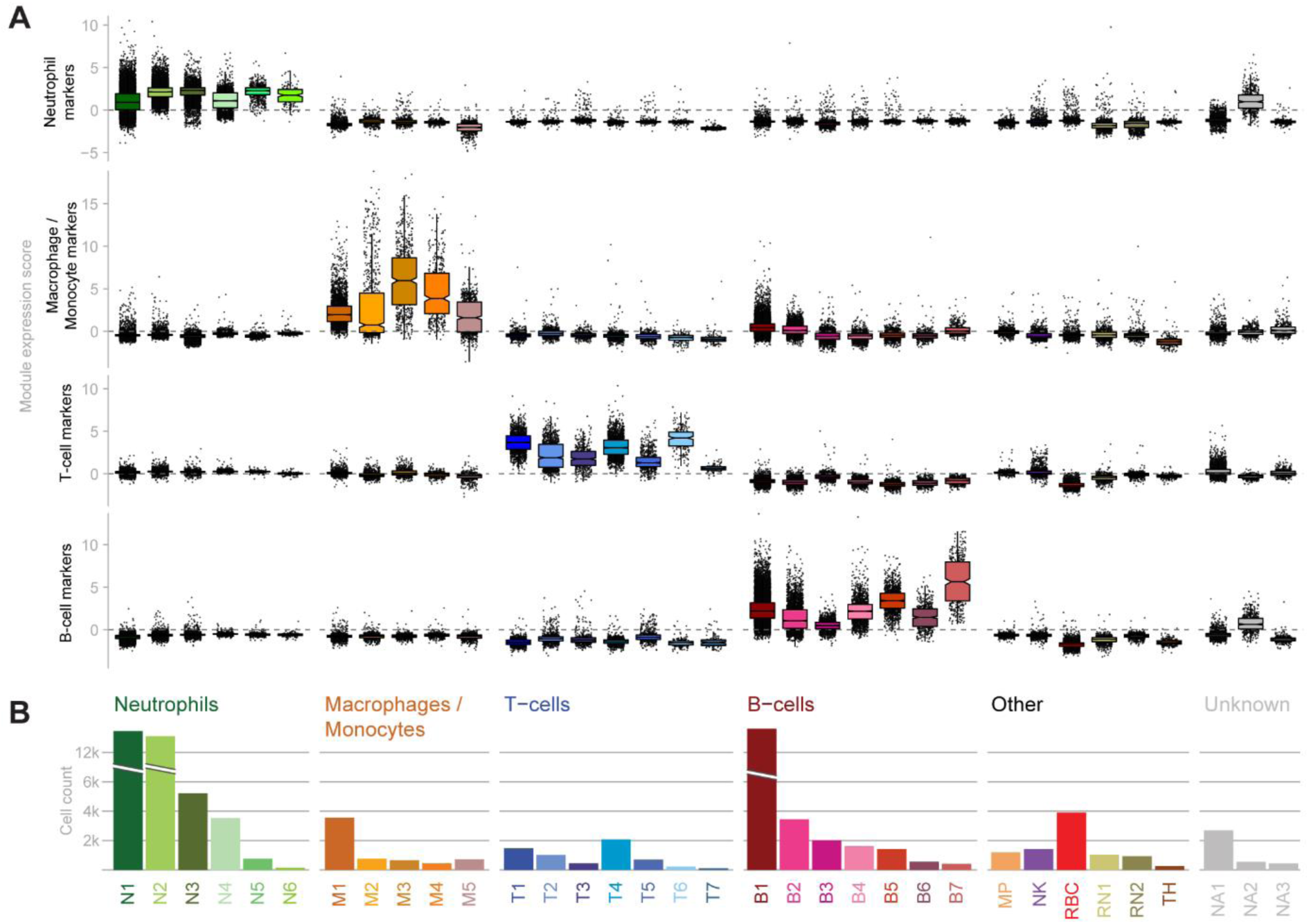
Cluster characteristics. (A) Module expression scores for Neutrophil markers (n=9), Monocyte / Macrophages (n=7), T-cells (n=22), and B-cells (n=12) across all clusters. **(B) Cluster identities and sizes.** Total number of cells assigned to each of the 34 clusters across all nine animals.

**Table 1.**
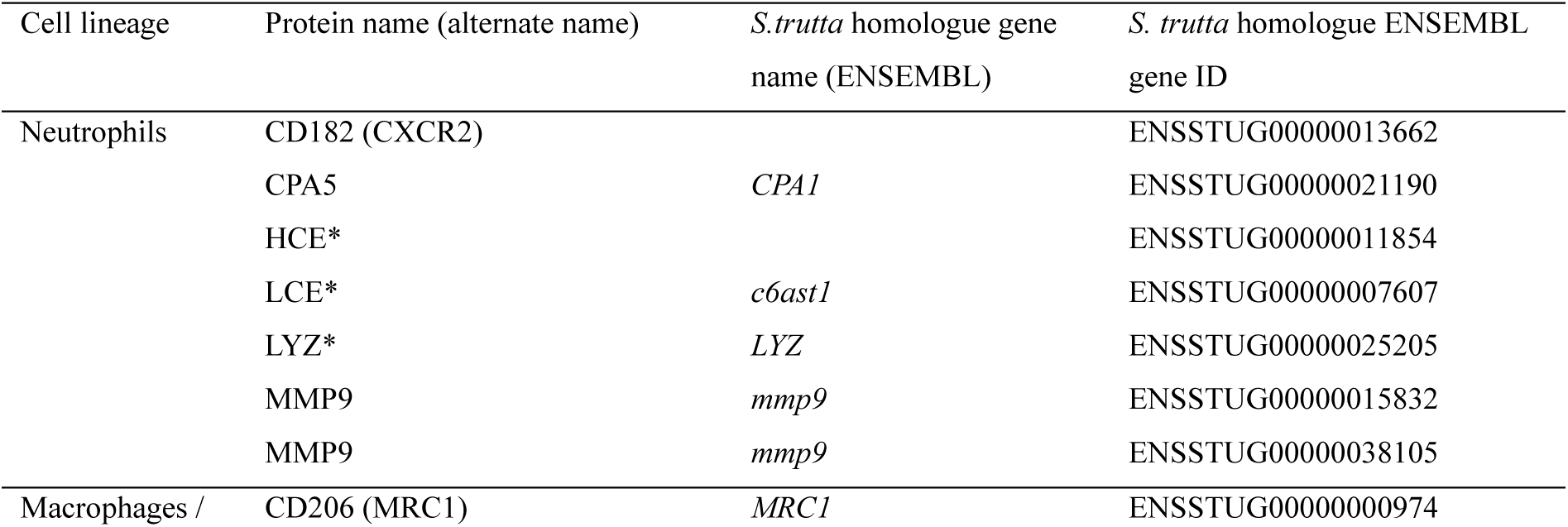
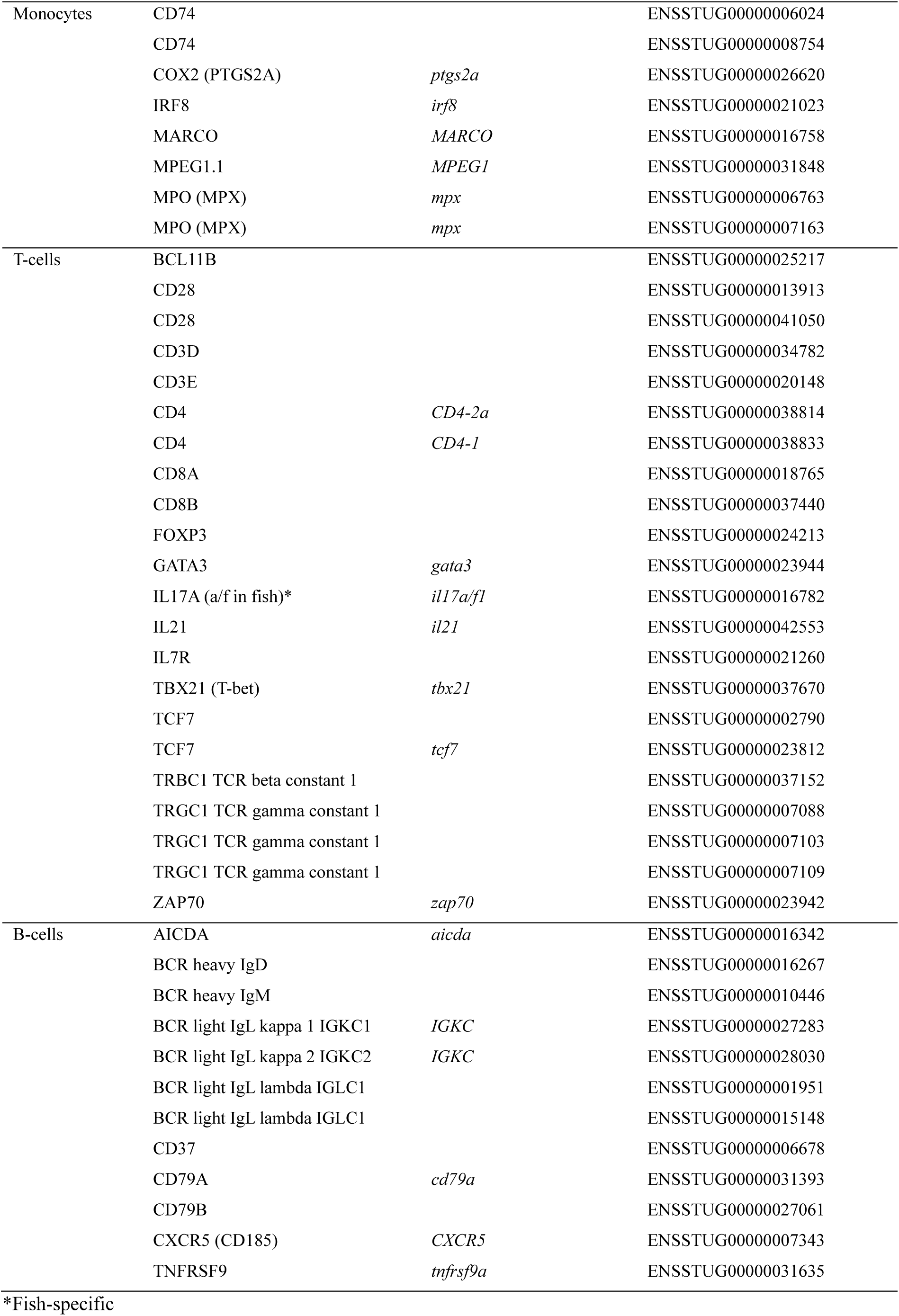
Prior markers of four major immune cell lineages whose S. trutta homologues were variably expressed in the scRNAseq dataset.

At a finer scale, putative cluster identities were inferred from genes that were overexpressed in the respective cluster at least 2-fold and in 60% of the cells (hereafter ‘cluster markers’; **Table S2**). Of 623 genes meeting these criteria, 600 were ENSEMBL genes and 23 were identified from our augmented annotation. 295 cluster markers were characteristic for a single cluster, and 328 cluster markers characterized more than one cluster. Prior markers and cluster markers together identified the putative cell type for 31 out of 34 clusters (80,166 out of 83,847 cells) (**Fig. 1B**). Two clusters were assigned to non-immune functions (RN1 and RN2, renal cells). Three remaining unidentified clusters (NA1, NA2 and NA3) were not assigned either due to lack of prior marker expression, lack of cluster marker expression in a high enough proportion of cells, or ambiguous expression patterns (e.g. markers of multiple cell types). Cluster NA2 in particular showed elevated expression of both B-cell and neutrophil markers and thus could not be assigned to either lineage with certainty.

### Proliferation markers identify several progenitor populations

Actively proliferating cell populations were identified by two prior markers of cell proliferation: *pcna* (Proliferating cell nuclear antigen; two homologues) and *mki67* (one homologue) and are found in neutrophil cluster N3, T-cell cluster T6 and B-cell cluster B4, within a subpopulation of macrophage cluster M1 and of putative myeloid progenitor cluster MP (**Fig. 3**). The identification as proliferating progenitor cells was further supported by expression of *H2AZ* (Histone variant H2AZ) and *pclaf* (PCNA clamp associated factor; **Fig. 3B**).

**Figure 3.**
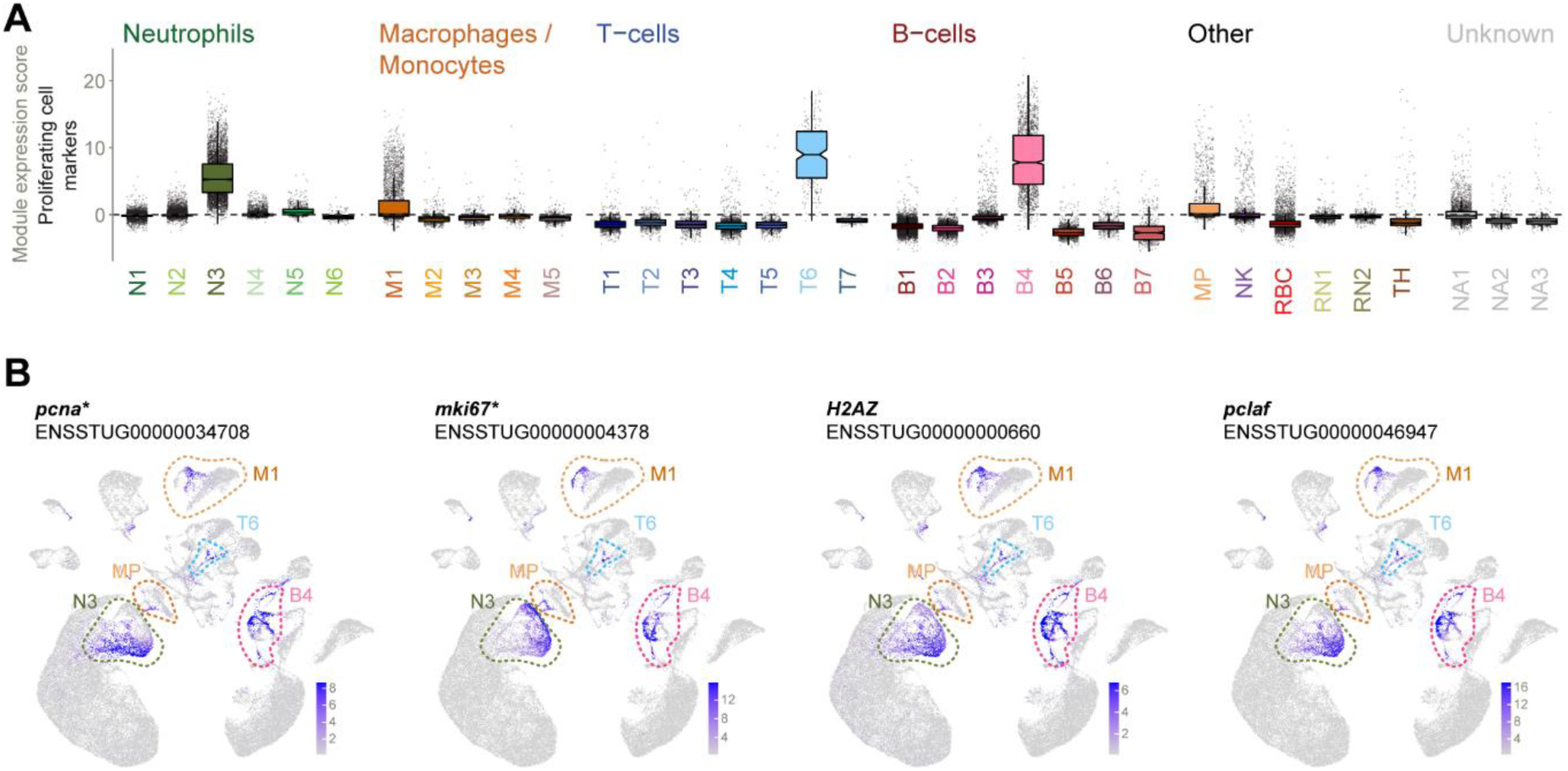
Proliferating cellS. (A) Module expression scores for proliferation markers (n=3) across all clusters. Clusters N3, M1, T6, B4 and MP contain substantial proliferating (sub)populations. **(B) UMAP projections** highlighting cells expressing two prior markers of proliferation (pcna and mki67), and two further genes (H2A.Z and PCNA caf) with similar expression patterns and cell cycle functions. Clusters showing high proportions of inferred proliferating cells are circled. Gene names are as given in the ENSEMBL annotation (without altering case). *Denotes gene is in a list of prior markers used to infer putative cell type or state.

### Neutrophils

In summary, we identify two major neutrophil subpopulations with specialized roles: one characterised by high expression of HCE1-like and another by abundant metallopeptidase gene expression. Overall, neutrophil clusters (N1 to N6, **Fig. 4A**) formed a large somewhat unstructured agglomeration, consistent with neutrophils being a) the most abundant immune cell type and b) a rather homogeneous population compared to other immune cell types. Neutrophil heterogeneity is however increasingly appreciated (Qu et al., 2023)), and indeed, we find that neutrophil transcriptional profiles are heterogeneous. Of 78 high abundance cluster markers (**Table S3**), none were common to all six subtypes (**Fig. 4B**).

**Figure 4.**
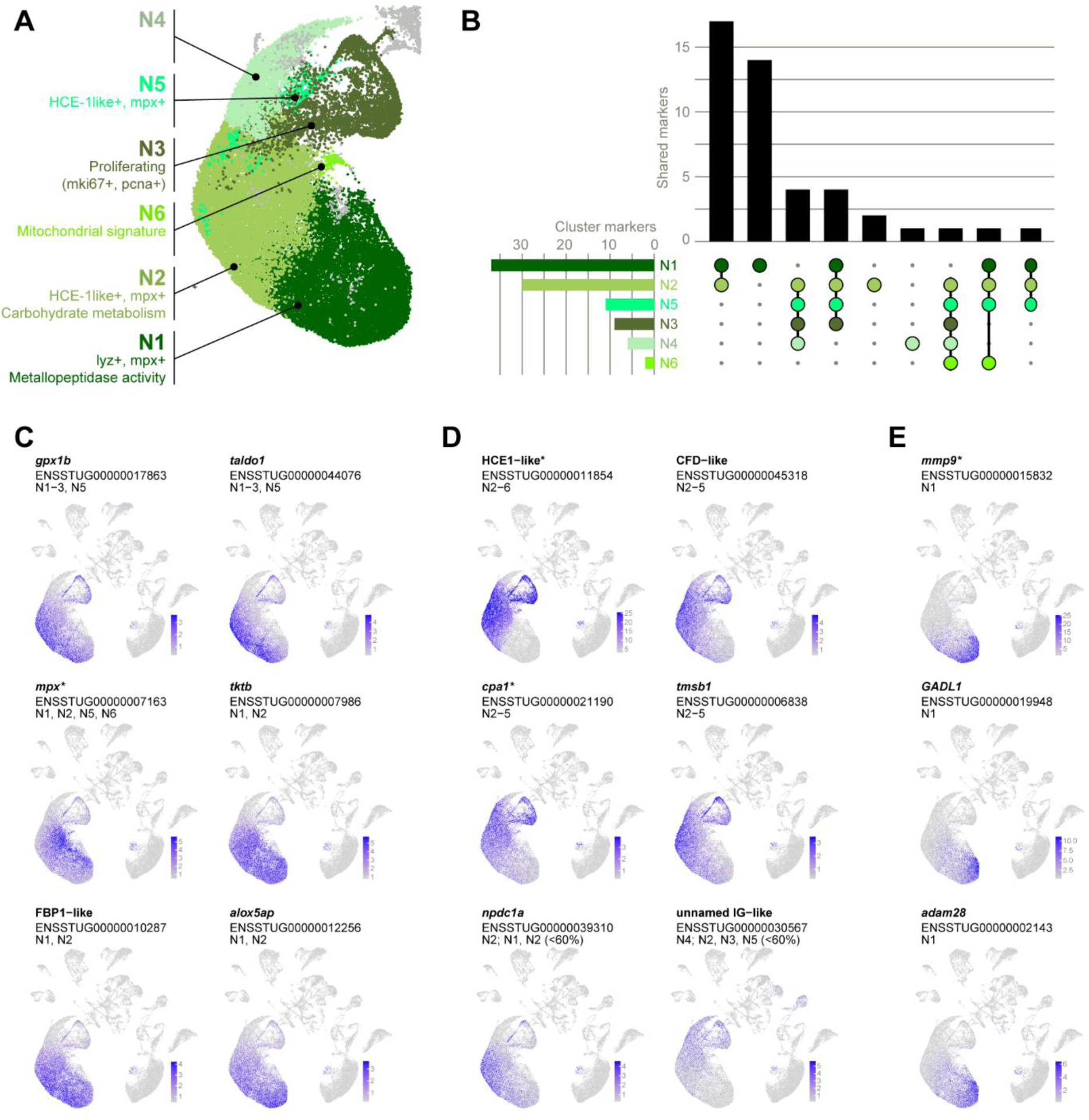
Neutrophils. (A) Distinguishing features of neutrophil clusters N1-N6. **(B) Unique vs shared high abundance cluster markers** (expressed in at least 60% of cells in the cluster). Markers shared with non-neutrophil clusters were excluded. **(C – E) Examples of cluster marker expression patterns.** Normalised expression values of markers **(C)** shared by the major Neutrophil clusters N1 and N2, **(D)** biased towards N2, and **(E)** expressed in N1. Gene names are as given in the ENSEMBL annotation (without altering case) except for entries with no gene names, in which case they are named according to protein homology (e.g. FBP1-like). Clusters for which the gene is a high abundance marker (expressed in at least 60% of cells) are listed. Where relevant, clusters for which the gene is a non-high abundance marker (expressed in < 60% of cells) are also indicated. *Denotes genes that are also prior markers.

Most neutrophils were collected in N1 and N2, and these two clusters also shared the largest number of markers, suggesting that markers of N1 and N2 can serve as a proxy for neutrophil fate. This set of “pan-neutrophil” markers includes homologues of human *taldo1*, *tktb* and FBP1 (**Fig. 4C**), which play a role in neutrophil responses to LPS exposure (Britt et al., 2022; Sadiku et al., 2021). Markers of N1 and N2 were generally enriched for the GO term ‘carbohydrate metabolic processes’, indicating a high energetic demand consistent with pro-inflammatory cells.

A few markers were more restricted in their expression, and able to differentiate between N1 and N2. A striking example of an N2 marker is HCE1-like (high choriolytic enzyme 1-like, **Fig. 4D**). HCE is responsible for breaking down the chorion during hatching of teleost fish (Nagasawa et al., 2022) but has more recently emerged as a marker of teleost granulocytes (Perdiguero et al., 2024; Wu et al., 2021). Other N2-biased markers include CFD1-like (complement-factor D-like), the human homologue of which inhibits leukocyte degranulation and thus prevents host tissue damage (Balke et al., 1995) and *cpa1*, a prior neutrophil marker and metallopeptidase.

N1 markers were consistently enzymes including the metalloproteases *mmp9, adam28*, and *mmp13*, and glutamate decarboxylase *GADL1* (**Fig. 4E**), and ‘metallopeptidase activity’ was a significantly enriched GO term amongst N1 markers. Lysozyme (*lyz*), previously identified as a neutrophil-specific marker in zebrafish (Harvie & Huttenlocher, 2015), was was also active in macrophages (see **Fig. S3**; in line with (Keshav et al., 1991)). Pseudotime analysis of neutrophil clusters (**Fig. S4A**) rooted at the putative myeloid progenitor cluster (MP, see **Fig. 8**) which implies that N1 cells are terminally differentiated.

N3 was identified as a pool of actively proliferating cells (see **Fig. 3**) while N4 and N5 lacked unique markers and may comprise intermediate or immature cells. N6 may represent dead or dying cells given a pattern of high mitochondrial gene expression (**Fig. S2**), fewer expressed genes, and low overall gene expression (Ilicic et al., 2016).

### Macrophages

Five clusters expressed prior markers of macrophages and / or monocytes and were termed M1 (a large cluster) and four small, fragmented clusters (**Fig. 5A**). Makrophages in brown trout are characterized by two prior markers, *MPEG1* and *irf8*, and two copies of *CD74*, which has been identified as a monocyte lineage marker in zebrafish (Athanasiadis et al., 2017). Another 25 high abundance cluster markers were unique to subsets of macrophages (**Table S4**), mostly to the largest cluster M1 (**Fig. 5B**). Antigen- presenting cell markers **(Fig. 5E**) consistent with canonical macrophage and dendritic cell function were broadly expressed, but beyond APC, only few markers were shared between individual macrophage clusters, suggesting functional heterogeneity.

**Figure 5.**
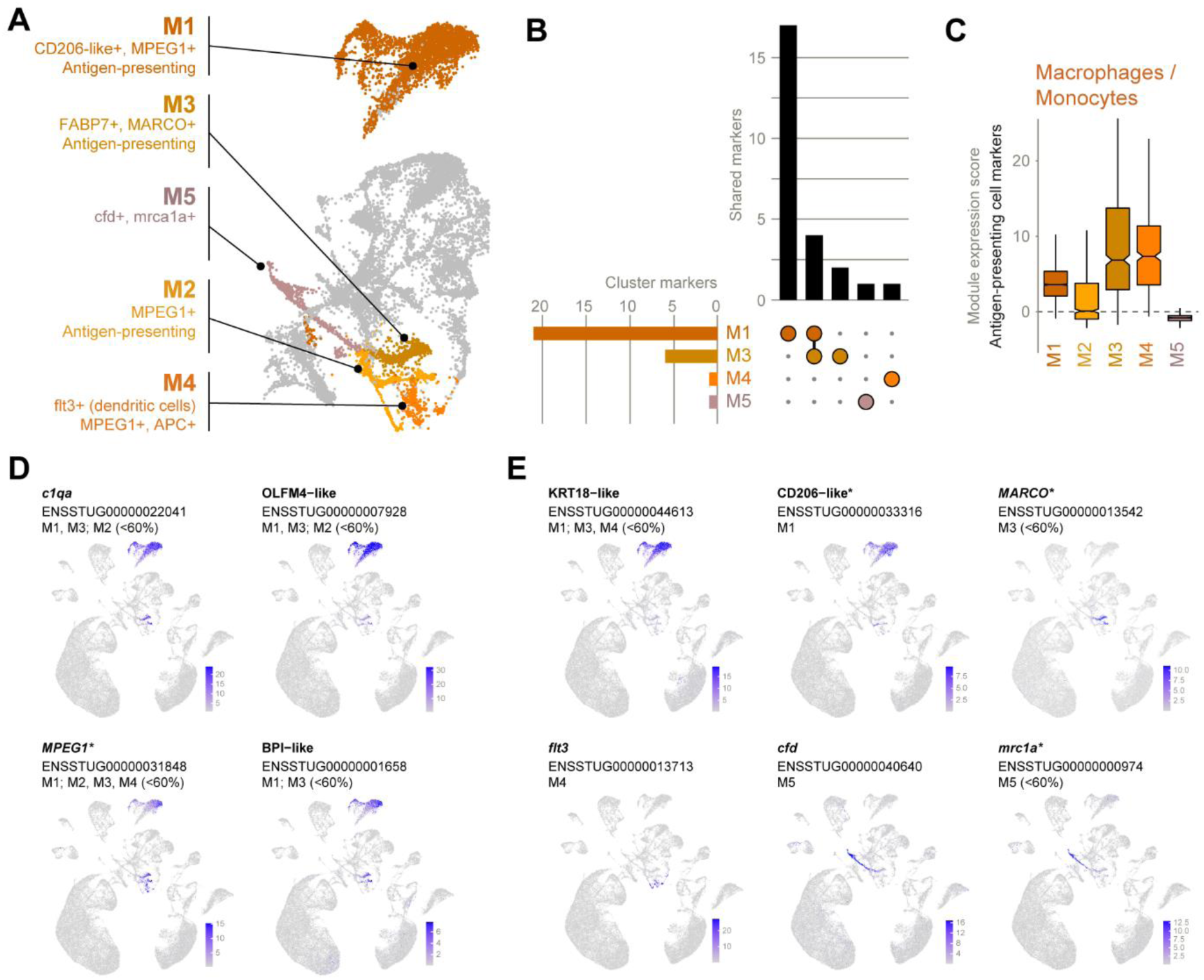
**Monocyte / macrophages**. **(A) Distinguishing features** of monocyte/macrophage clusters N1-N6. **(B) Unique vs shared high abundance cluster markers** (expressed in at least 60% of cells in the cluster). Markers shared with non-macrophage clusters were excluded. **(C) Antigen-presenting cells.** Expression of a gene module comprised of CD40, CD74 (2 homologues), MHCII beta−like, and MPEG1. **(D - E) Examples of cluster marker expression patterns.** Normalised expression values of markers **(C)** shared by multiple clusters and **(D)** expressed principally in one cluster. Gene names are as given in the ENSEMBL annotation (without altering case) except for entries with no gene names, in which case they are named according to protein homology (e.g. BPI-like). Clusters for which the gene is a high abundance marker (expressed in at least 60% of cells) are listed. Where relevant, clusters for which the gene is a non-high abundance marker (expressed in < 60% of cells) are also indicated. *Denotes genes that are also prior markers.

M1, the largest and most distinct cluster with the highest number of unique high abundance markers was characterized by CD206-like (macrophage mannose receptor 1 homologue) and FCGR1-like (**Fig. 5D**). In mammals, both have roles in phagocytosis (Azad et al., 2014)(Swanson et al., 1999). The smaller, fragmented clusters featured fewer expressed genes, lower overall expression and higher mitochondrial gene expression (**Fig. S2**), suggesting that they are comprised of or contained dead or dying cells.

M3 shared four high-abundance markers with M1: three paralogues of complement component 1q (*c1qa*, *c1qb*, and *c1qc*) and OLFM4-like (**Fig. 5C**). Similarly, GO term enrichment analysis showed markers of M1 and M3 to be enriched for ‘immune response’ and ‘antigen processing and presentation’ (**Table S7**).Unique M3 markers include *FAPB7* (fatty acid binding protein 7) and *epdl1* (ependymin). Fatty acid metabolism is important in macrophages to produce inflammatory mediators (Batista-Gonzalez et al., 2020), while ependymins are a class of purportedly teleost-specific glycoproteins found in cerebrospinal fluid and involved in the central nervous system (McDougall et al., 2018). Around half of M3 cells also expressed the classical macrophage marker *MARCO*.

M4 expressed the unique marker *flt3*. The murine homologue is a major dendritic cell growth factor (Karsunky et al., 2003), thus implying a dendritic fate for M3. Although dendritic cells are thought to derive from a distinct cell lineage from monocytes, may be functionally similar and thus may resemble monocytes transcriptionally (Guilliams et al., 2014). Uniquely, M5 contained cells expressing classical macrophage marker *mrc1a* (<60% of cells), despite not showing elevated Antigen-Presenting Cell (APC) marker expression (**Fig. 5E**). Generally, the relative scarcity of high abundance markers compared to genes expressed in subpopulations of clusters (25 high-abundance markers compared to 75 non-high-abundance markers after excluding markers of other clusters) suggests that the level of clustering in this study does not fully grasp macrophage heterogeneity.

### T-cells

Seven clusters were identified as T-cells based on prior markers and designated T1-7 (**Fig. 6A**). Interestingly, cluster T7 was represented almost exclusively by fish from the wild-caught group (**Fig. S5**). Given the potential of confounding of markers by genetic, environmental, or genotype-by-environment effects, we focused on clusters T1-T6 in our characterisation of T-cell subpopulations.

**Figure 6.**
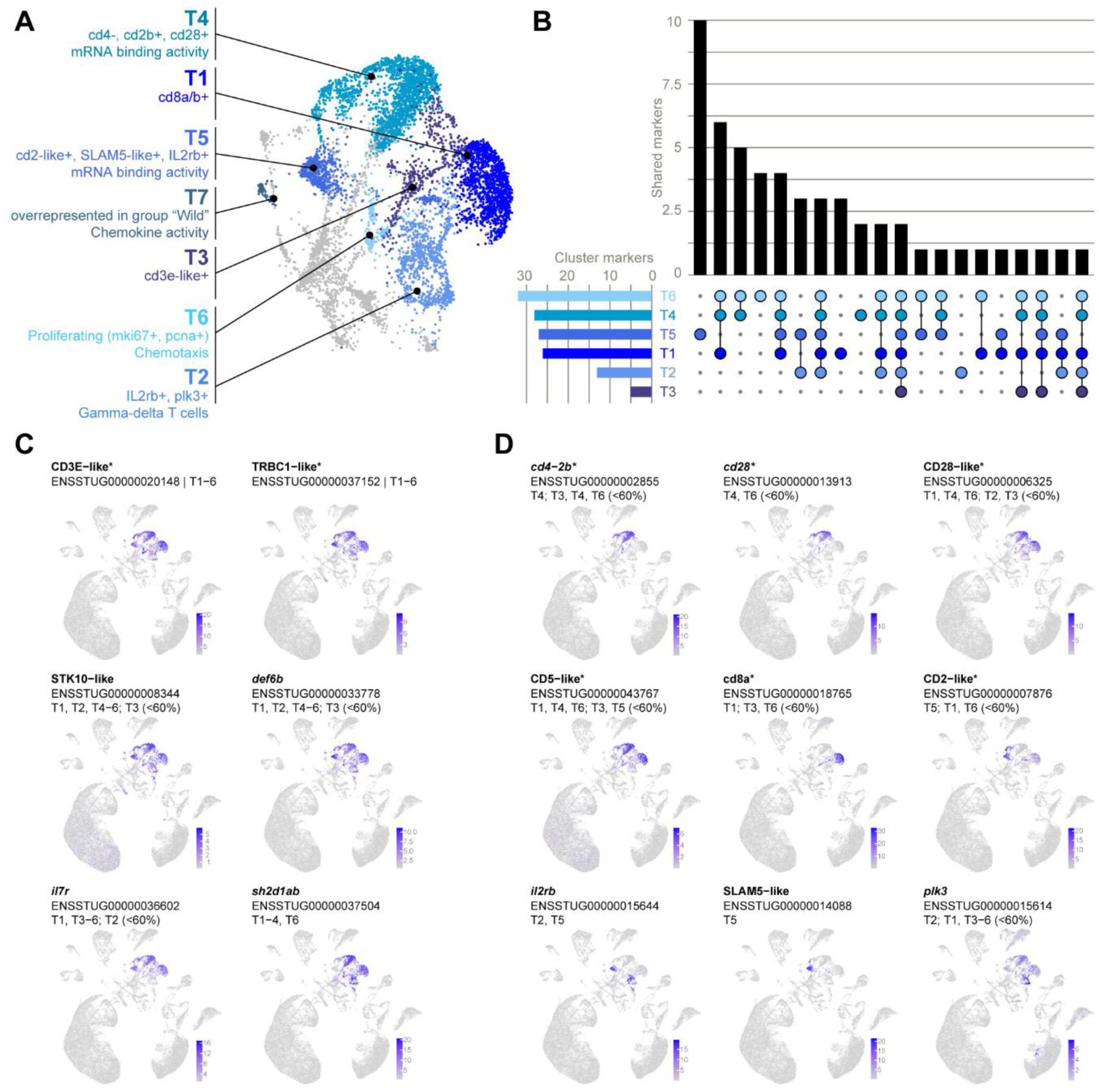
T-cells. (A) Distinguishing features of T-cell clusters T1-T7. **(B) Unique vs shared high abundance cluster markers** (expressed in at least 60% of cells in the cluster). Markers shared with non-T-cell clusters were excluded. **(C - D)**: UMAP projections showing **(C)** shared markers expressed in multiple T-cell clusters, **(D)** unique markers expressed principally in one T-cell cluster. **Examples of cluster marker expression patterns.** Normalised expression values of markers **(C)** shared by multiple clusters and **(D)** expressed principally in one cluster. Gene names are as given in the ENSEMBL annotation (without altering case) except for entries with no gene names, in which case they are named according to protein homology (e.g. SLAMS-like). Clusters for which the gene is a high abundance marker (expressed in at least 60% of cells) are listed. Where relevant, clusters for which the gene is a non-high abundance marker (expressed in < 60% of cells) are also indicated. *Denotes genes that are also prior markers.

Of 53 genes identified as T-cell markers (**Fig. 6B; Table S5**), only two can be considered ‘pan’ T-cell markers (**Fig. 6C**): CD3E-like and TRBC1−like (TCR beta constant 1-like), both of which were also prior markers and represent well-characterised T-cell surface receptors. A further five markers were shared by five clusters: STK10-like, *def6b*, *il7r* (interleukin 7 receptor), *sh2d1ab*, and *fynb*. STK10 is a putative immunoregulator expressed in human T-cells (but also in dentritic cells and NK cells; (Bi et al., 2022)). Human DEF6 is important for T-cell homeostasis (Serwas et al., 2019), while human SH2D1A is involved with SLAM-signaling (important for T-cell proliferation) and indeed has known interactions with FYN (Chan et al., 2003).

T1, T4, and T5 could be largely distinguished by expressed homologues of canonical T-cell markers, namely CD2, CD4, CD5, and two homologues of each of CD8 and CD28 (**Fig. 6D**; only one CD8 homologue shown). T1 represents CD8+ T-cells, as it uniquely expressed two homologues of CD8 in high abundance (*cd8a* and *cd8b*; latter not shown in Fig. 6B, but exhibited identical expression pattern to *cd8a* indicating its expression as a heterodimer), a surface protein characteristic of memory and cytotoxic T-cells (Macintyre et al., 2011). Interestingly, T1 expressed only one of two CD28 homologues, suggesting some degree of functional divergence. T1 markers were enriched for the GO term ‘integrin complex and indeed integrins are important for T-cell activation by promoting stable contact with antigen-presenting cells (Burbach et al., 2007). Meanwhile, T4 represents CD4+ cells, as it was distinguished by expression of both CD28 homologues along with the T-helper cell marker *cd4-2b*.

T5 was the most distinct T-cell cluster with 10 high abundance markers not shared by other clusters. It was characterised by expression of a homologue of canonical T-cell marker CD2 along with SLAM5- like (human homologue also known as CD84) and *il2rb*, which are markers of regulatory T-cells (Chinen et al., 2016; Cuenca et al., 2019). T2 expressed only one unique marker expressed in high abundance (likely reflecting a cluster composed of more than one distinct subpopulation which were not completely resolved) but expressed *plk3* in combination with *ilr2b*, the latter suggesting this cluster also contained regulatory T-cells. T2 also uniquely contained cells expressing two unnamed homologues of TRGC1 (TCR gamma constant 1; **Fig. S6**), suggesting that another subpopulation of T2 represents the less common gamma-delta T-cells, supporting the previous sequenced-based notion that these cells exist in salmonids (Yazawa et al., 2008).

T6 markers were consistent with proliferating cells (**Fig. 3**), while T3 did not express any unique high abundance markers and showed signatures of dead or dying cells (high mitochondrial gene expression, few expressed genes and low overall expression; **Fig. S2**).

### B-cells

Seven B-cell clusters were clearly separated on the UMAP (**Fig. 7A**). After excluding markers of non-B-cell clusters, 113 genes were identified as B-cell markers (**Fig. 7B; Table S6**). Several largely shared high-abundance markers were identified, but all seven clusters also possessed at least one unique marker (**Fig. 7B**). A marker shared at least by five clusters is *cd79a* (**Fig. 7C**). *cd79b* showed similar expression patterns, but was less expressed in the larger clusters B1 and B2. Other ‘pan’ B-cell markers included genes whose mammalian homologues have been demonstrated to be involved in B-cell proliferation or differentiation: *mef2cb* (Wilker et al., 2008), *hipk2* (Guerra et al., 2012), and one homologue of EBP41 (Liang et al., 2020), as well as canonical B-cell marker homologue CD37-like and at least six immunoglobulin-like genes (e.g. first two panels of **Fig. 7E**). In addition, B-cell markers included several genes homologous to Ig proteins, some lacking annotation but containing predicted IG domains (see **Table S6**). These tended to show variable expression between and within clusters (**Fig. 7E**) and included an IGLL1-like gene which was uniquely overexpressed in B3.

**Figure 7.**
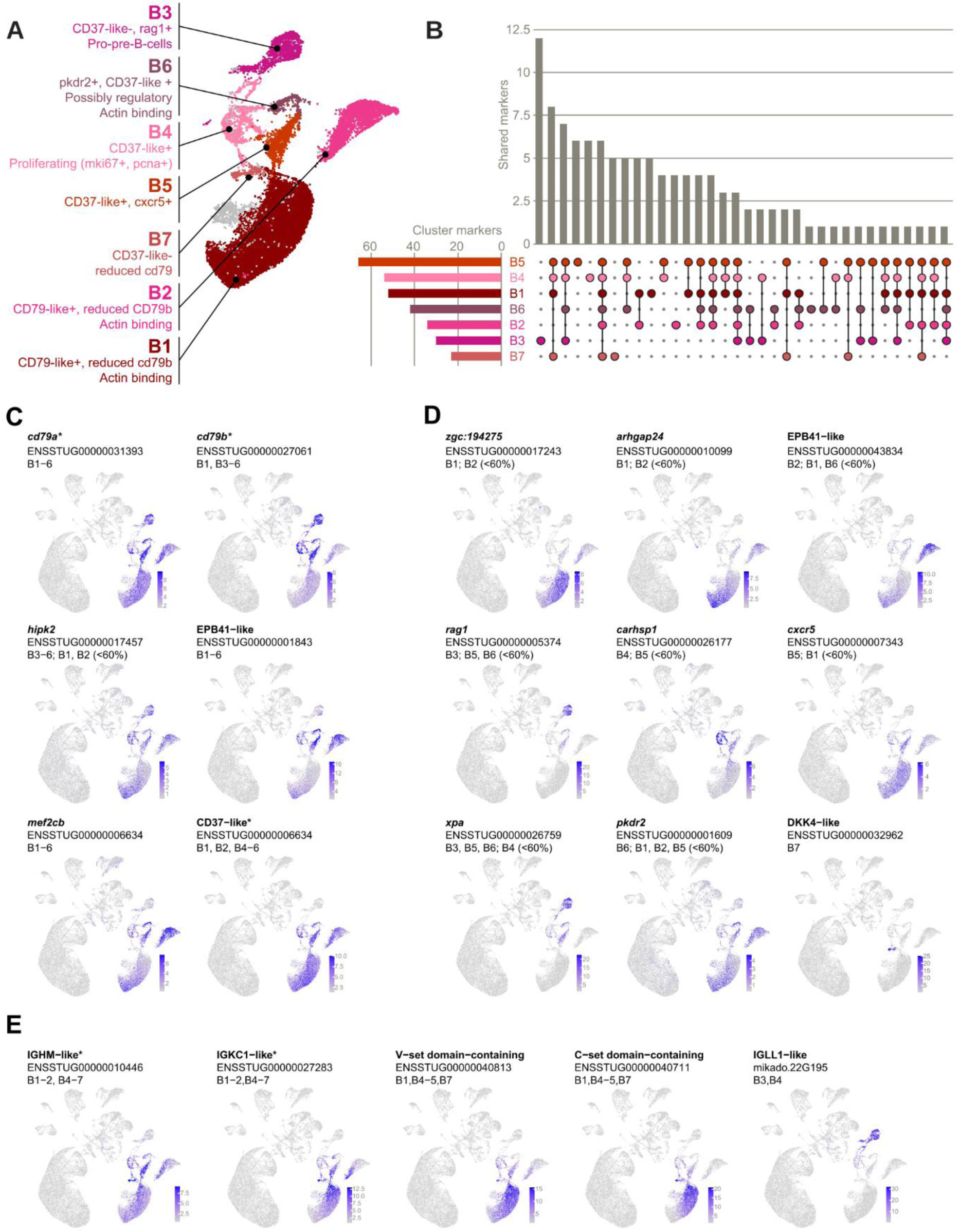
**B-cells**. **(A) Distinguishing features** of B-cell clusters B1-B6. **(B) Unique vs shared high abundance cluster markers** (expressed in at least 60% of cells in the cluster). Markers shared with non-T-cell clusters were excluded. **(C – E) Examples of cluster marker expression patterns.** Normalised expression values of markers **(C)** shared by multiple clusters, **(D)** expressed principally in one cluster and **(E)** immunoglobulin homologues or markers containing Ig domains. Gene names are as given in the ENSEMBL annotation (without altering case) except for entries with no gene names, in which case they are named according to protein homology (e.g. IGLL- like). Clusters for which the gene is a high abundance marker (expressed in at least 60% of cells) are listed. Where relevant, clusters for which the gene is a non-high abundance marker (expressed in < 60% of cells) are also indicated. *Denotes genes that are also prior markers.

B1 and B2 (putatively mature B-cells) display overlapping expression patterns. High-abundance markers of B1 are often also highly expressed in B2 (though not necessarily in high abundance), and *vice versa* (**Fig. 7D**). *cd79b* was only weakly expressed in both, which in zebrafish is a sign for differentiation to the Ig-secreting stage (Liu et al., 2017). *arhgap24*, which is enriched in naïve and memory B-cells in humans (Human Protein Atlas version 24; https://www.proteinatlas.org/ENSG00000138639-ARHGAP24/single+cell (Uhlen et al., 2019)) and inhibits cell proliferation (L. Wang et al., 2019), marked both B1 and B2. Ig-like genes were also highly expressed in B1 and B2 (**Fig. 7E**), and B1 and B2 markers were enriched for the GO term ‘actin binding’; indeed, reassembly of the B-cell actin cytoskeleton allows expansion of the cell surface to increase contact with antigen-presenting cells (Song et al., 2013).

Most specific to B1 was *zgc:194275*, a homologue of a zebrafish B-cell marker (C. Hu et al., 2024) also known as PLAAT1, which regulates antiviral responses in zebrafish (Zhao et al., 2022). Pseudotime analysis of B-cell clusters with B3 assigned as the starting point (**Fig. S4B**) supported the designation of B1 as terminally differentiated B-cells.

Specific B2 markers include the second homologue of the pan-B-cell marker EBP4, implying functional divergence, and a previously unannotated gene (mikado.19G1247, not shown) with homologies to NHSL2, a protein involved in cell migration – a process critical for patrolling immune cells (Y. Wang et al., 2023).

B3 unique markers support a putative pre- or pro-B-cell identity. Most notably, B3 highly expressed *rag1* (recombination activating gene 1), a gene well known for its crucial role in B-cell development (Kuo & Schlissel, 2009), see also (Novoa et al., 2019). Other markers of B3 shown to have roles in B-cell or pre-B-cell development include a*kap12* (Kartal-Kaess et al., 2012), *itga5* (Y. Hu et al., 2016), and *tek* (Reth & Nielsen, 2014), and markers shared with B4 were enriched for the GO term ‘structural constituent of chromatin’ suggestive of chromatin remodeling and differentiation.

B4 was a highly proliferative cluster as evidenced by expression of proliferation markers (see **Fig. 3**). B5 was charactized by *cxcr5*, a marker associated with B-cell migration (Förster et al., 1996) and with B helper T cells (Breitfeld et al., 2000). B5 and B6 shared *xpa* (associated with DNA repair; (Kim et al., 2023) as a marker with the pro/pre-B-cell cluster B3, suggestive of an intermediate stage between pro/pre-B-cells and mature B-cells.

B6 represents putative regulatory B-cells based on expression of *pkdr2* (Michaud et al., 2022). Also, B6 markers were enriched for actin binding similarly to B1 and B2, suggesting similar interactions with antigen-presenting cells.

Finally, B7 was a small cluster of seemingly highly differentiated cells as indicated by lack of expression of both CD79 genes (Liu et al., 2017) and intense expression of Ig-like genes (see **Fig. 7E**). Unique high-abundance markers of B7 included DKK4-like, a similar gene to which in mammals, DKK3, is a modulator of B-cell fate and function (Ludwig et al., 2015).

### Other distinct cell types are resolved by marker genes

Natural killer-like cells (cluster NK) were identified by prior markers such as *prf1* (perforin) and characterized by four unique cluster markers which also included a previously unannotated homologue of NK-lysin (**Fig. 8A**).

A small but highly distinct cluster (41 unique markers) corresponds to thrombocytes (cluster TH) according to expression of prior markers such as *mpl* (thrombopoietin receptor-like) and *thbs1b* (thrombospondin 1-like) (**Fig 8B**), both of which have mammalian homologues involved in platelet formation and / or aggregation (Hitchcock et al., 2021; Leung, 1984).

Red blood cells (cluster RBC) were identified based on expression of hemoglobin homologues such as *hbaa2* (**Fig. 8C**). Eleven genes were markers uniquely of the RBC cluster, including *alas2* (5- aminolevulinate synthase, erythroid-specific, mitochondrial-like), thus comprising markers of mature RBCs. Hemoglobin homologues were also sporadically, but highly expressed in the central unclassified cluster NA1, suggesting it contained erythroid progenitors.

Another small cluster was identified as putatively comprising myeloid progenitor cells (cluster MP), whose unique markers included *kita* and *egr1* (**Fig. 8D**), homologues of which are heavily implicated in early myeloid cell differentiation (Ashman et al., 1999; Kulkarni, 2022). Among other markers of MP were three *fos* genes and a homologue of transcription factor JUND which, along with EGR1 are all implicated in early myeloid function (Yuan et al., 2007).

Finally, two clusters with similar expression patterns were putatively identified as renal (kidney) cells (clusters RN1 and RN2) based on markers such as *ly97.3* (CD59-like; **Fig. 8E**), *si:dkey-194e6.1* (collectrin-like) and *gstr* (glutathione S-transferase A-like), mammalian homologues of which are highly expressed in cells of the kidney (Harrison et al., 1989; Nangaku et al., 1998; Zhang et al., 2001). RN1 and RN2 were also the only clusters showing appreciable expression of *gapdh* (**Fig. 8E**).

### Functional divergence of gene pairs in the immune system

Several marker genes used to identify clusters were present in duplicate, and displayed differential expression between clusters, for example CD28 (**Fig. 6D**) and EBP41 (**Fig. 7D**). We therefore systematically investigated the expression patterns of duplicate gene pairs arising from the salmonid-specific ‘SSR4’ whole genome duplication event (hereafter ‘ohnologues’), based on the assumption that differential expression could signify subfunctionalisation (i.e. division of labour) or neofunctionalisation (i.e. one copy gains a new function). First, we identified 175 putative ohnolog pairs amongst the variably expressed genes by querying the 2000 variably expressed genes for homologues in the northern pike (*E. Lucius*).

From 176 ohnologue pairs (**Table S8**), 87 show identical or near-identical expression patterns across clusters (see examples in **Fig. 9A**), suggesting that both have retained the same function or have functionally diverged only on a subcellular level. 68 gene pairs display ‘gain or loss’ patterns, where ohnologues have similar expression but one ohnologue is missing from certain clusters (see examples in **Fig. 9B**), which may suggest either subfunctionalisation or a transition to neofunctionalization. 6 gene pairs display ‘partially contradicting’ patterns, where each ohnologue is to some extent expressed in mutually exclusive patterns while retaining some overlap, suggesting the beginning of neofunctionalization. 15 gene pairs display ‘contradicting’ patterns, where ohnologues display entirely mutually exclusive patterns, suggesting neofunctionalisation (see examples in **Fig. 9C**).

**Figure 8.**
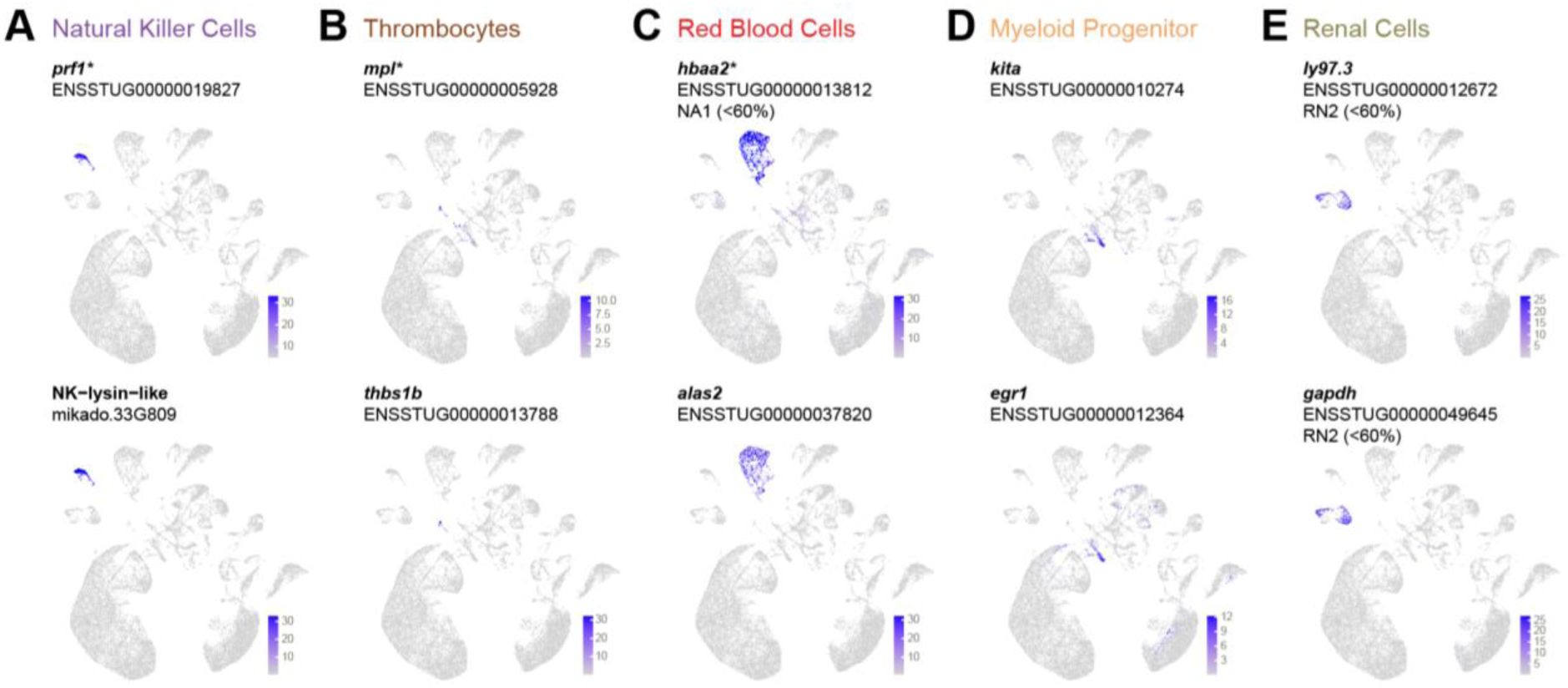
**Other cell types**. **Examples of cluster marker expression patterns.** Normalised expression values of markers for **(A)** natural killer cells, **(B)** thrombocytes, **(C)** red blood cells, **(D)** myeloid progenitor cells, and **(E)** renal cells. Gene names are as given in the ENSEMBL annotation (without altering case) except for entries with no gene names, in which case they are named according to protein homology (e.g. NK-lysin-like). Where relevant, clusters for which the gene is a non-high abundance marker (expressed in < 60% of cells) are also indicated. *Denotes genes that are also prior markers.

**Figure 9.**
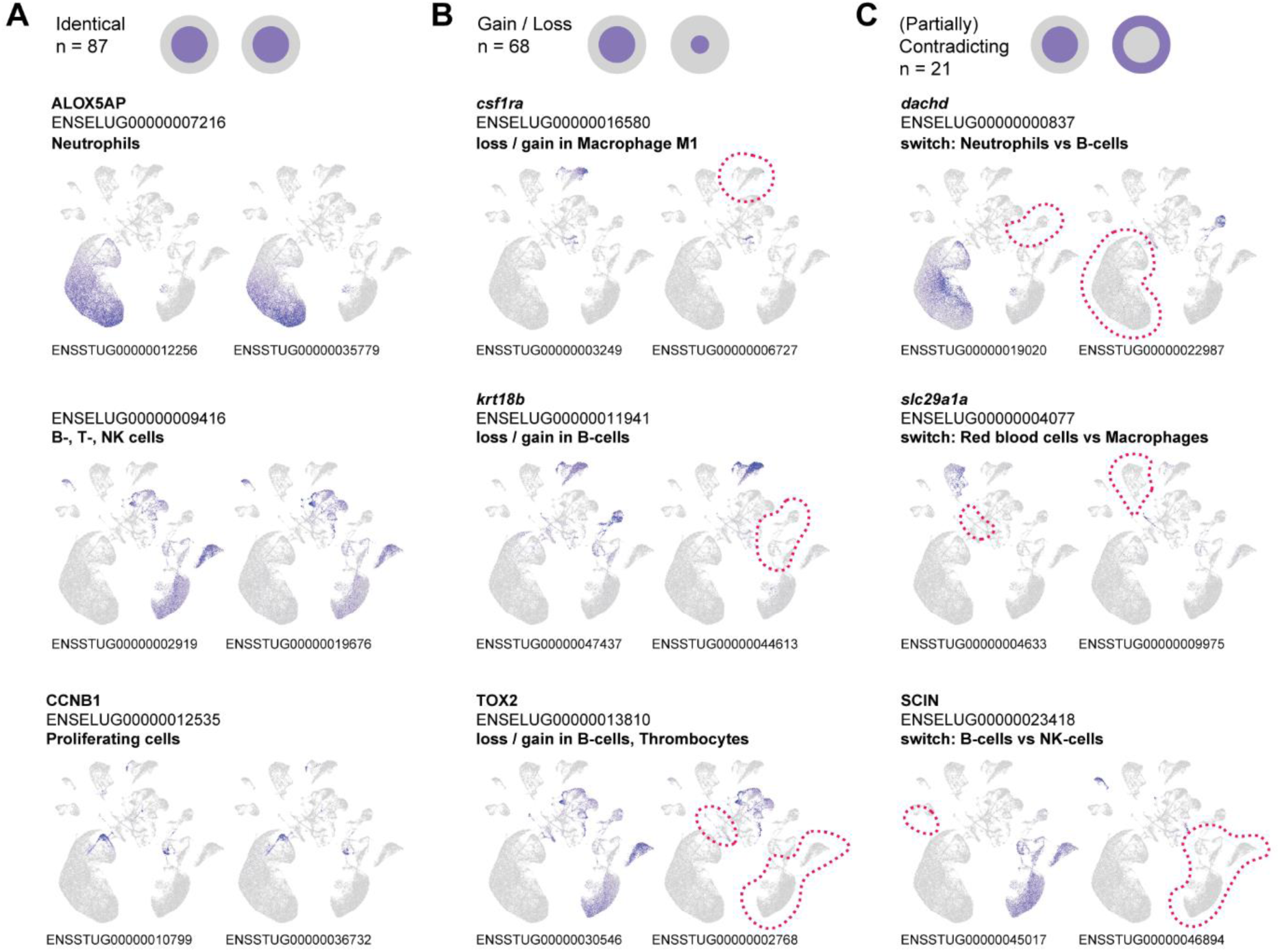
Ohnolog neofunctionalisation. Ohnologue pairs with **(A)** identical expression patterns, **(B)** one ohnologue missing from certain clusters **(C)** asymmetric expression patterns. ENSELUGXXXX: ENSEMBL gene ID and gene names of the E. lucius single copy homolog. Red dotted lines: clusters that lost expression compared to the other ohnologue.

A striking example of contradictory expression patterns between ohnologes is *dachd.* The Drosophila Dach transcription factor (*dachd* homologue) is a master developmental regulator, forming complexes with chromatin to induce differentiation (Davis & Rebay, 2016). In brown trout, one ohnologue was expressed widely in neutrophils and the other in pre-B-cells. The ohnologs of *SCIN*, an actin-severing protein (Ghoshdastider et al., 2013), were expressed in B-cells versus NK-cells (**Fig. 9C**). Plots of all examined ohnolog pairs can be found in **Supplementary File 1**.

### Environment of current or prior generations affects immune cell transcriptomes

Our data shows that rearing environment impacts immune gene expression and even the presence of immune cell clusters. Briefly, ‘wild’ fish (N =3) were captured directly from a local stream in the canton of St. Gallen, Switzerland, and had an intact fat fin indicating that they were born and raised in the wild. The ‘farm’ fish (N =3) were derived from at least two generations of rearing in an aquaculture facility, while ‘mix’ fish (N =3). Interestingly, and as noted earlier, T-cell cluster T7 was represented almost completely exclusively by fish from the ‘wild’ group (Fig. S5) and was the only such cluster to display an origin-specific abundance pattern, suggesting distinct subset that farmed fish fail to produce.

To characterise the influence of multigenerational and single-generation hatchery rearing, we selected a cluster from each of the four main lineages (N1, M1, T5 and B3 for neutrophils, monocytes / macrophages, T-cells, and B-cells, respectively) and performed between-group differential expression analyses using the pseudobulk approach (Murphy et al., 2022).

Of the selected cell types, macrophages (represented by M1) showed the most abundant differences in gene expression between groups (235 DEGs at FDR < 0.05, of which 157 were DE in mix and 78 in farm; **Table S9**), followed by neutrophils (represented by N1; 178 DEGs at FDR < 0.05, of which 95 were DE in mix and 83 in farm; **Table S10**) and B-cells (represented by B3; 121 DEGs at FDR < 0.05, of which 65 were DE in mix and 56 in farm; **Table S11**). PCA plots of all three showed separation between rearing conditions along PC1 and / or PC2 (**Fig. 10**). T-cells (represented by T5) meanwhile showed more modest differences (74 DEGs at FDR < 0.05, of which 35 were DE in mix and 39 in farm; **Table S12**) and less clear sample separation on PCA (**Fig. S7**).

**Figure 10.**
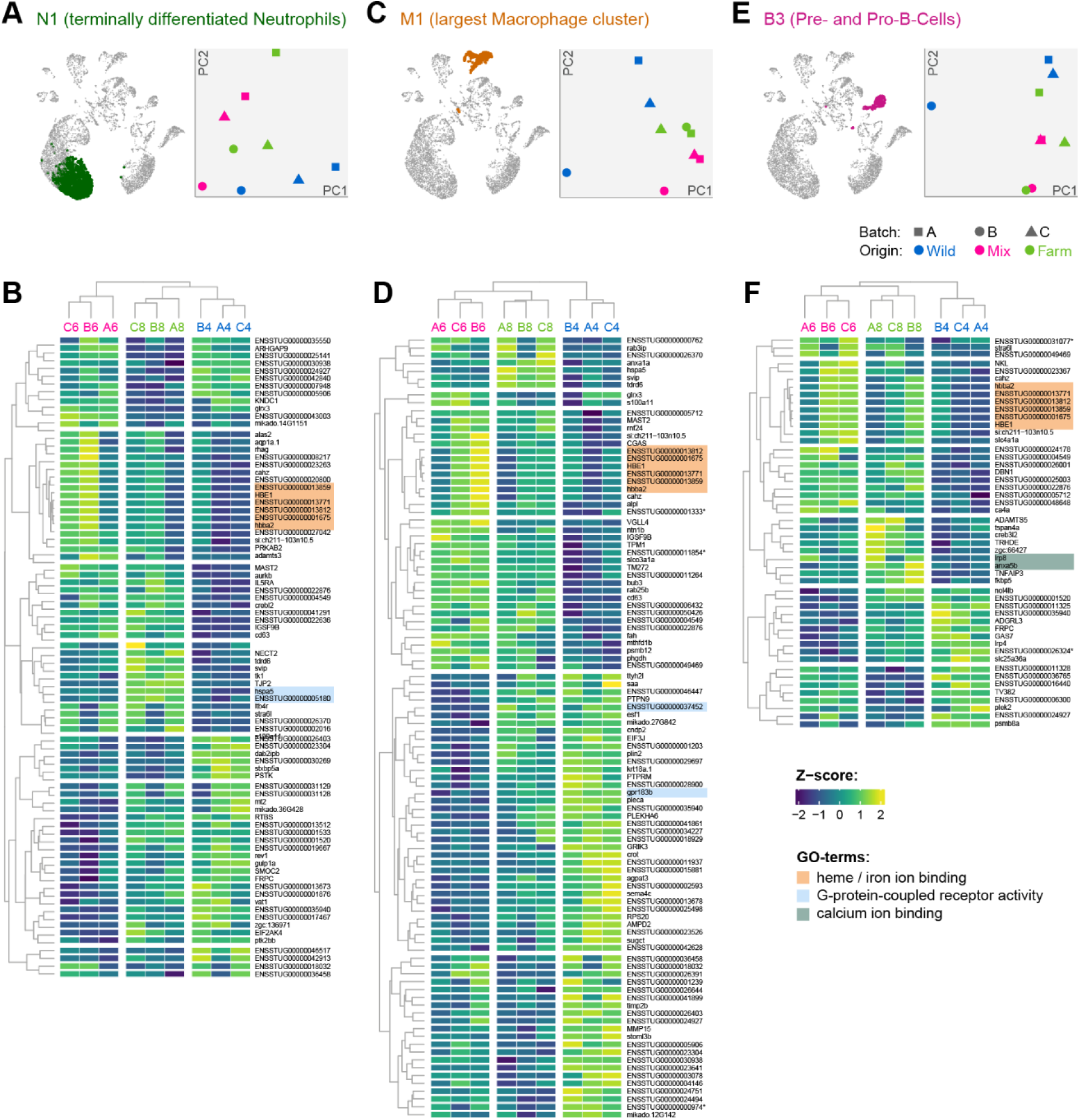
Impact of raising environment. (A, C, **E)**: Terminally differentiated Neutrophils, Macrophages, and Pre-/Pro-B-cells highlighted on UMAP plot, and PCA plot of all samples stratified by raising environment (point color) and experimental batch (point shape). **(B, D, F)**: Genes differentially expressed (FDR < 0.01) between ‘farm’ and/or ‘mix’ compared to the ‘wild’ group. Dendrograms represent hierarchical clustering of genes and samples according to the Z-scores of the Log2(CPM) values. Cell color indicates low (blue) versus high (yellow) z-score. Sample IDs at the top of each heatmap indicate raising environment (wild = blue, farm = pink, mix = green). Genes are indicated by gene name where available from the ENSEMBL database, or by ENSEMBL gene ID. Genes corresponding to GO terms enriched amongst DEGs are highlighted (red = ‘heme binding’ or ‘iron ion transport’, blue = ‘G protein-coupled receptor activity’, green = ‘calcium ion binding’). Note that genes DE with FDR > 0.01 are not shown but were used for GO term analysis.

To explore potential functional implications this rearing-induced differences, we ran GO term enrichment analysis using the GO domain ‘molecular function’, which we considered more appropriate than biological processes or cellular component given that the cell types had already been inferred. Surprisingly, similar GO terms emerged for neutrophils, macrophages, and B-cells (**Table S13**), suggesting a pan-cell type effect of early life experiences on the immune system. ‘heme binding’ and ‘iron ion binding’ were upregulated in the mix group in all three aforementioned clusters, ‘G protein-coupled receptor activity’ was upregulated in the farm group in Neutrophils and B-cells, and downregulated in the mix group in macrophages. In B-cells, ‘calcium ion binding’ was additionally enriched amongst genes upregulated in the farm group. No enriched GO terms were detected for genes differentially expressed in T5.

It is worth noting that genes involved in ‘heme binding’ and ‘iron ion binding’, comprising putative hemoglobin homologues, exhibited a batch effect in that they were consistently downregulated in batch ‘A’, regardless of origin.

## Discussion

Our data suggests a conserved immune system setup for brown trout, given that expression modules derived from other species’ immune cell markers readily identified core cell types. This is noteworthy because an increasing number of fish immune systems have been analyzed in detail, e.g. through single cell RNA sequencing (Fuess & Bolnick, 2023; Hu et al., 2024; Chen et al., 2022; Wang et al., 2024), and some were found to notably differ from the vertebrate core configuration (Guslund et al., 2020; Guslund et al., 2022). Broadly, our results are therefore in agreement with previous scRNAseq studies of salmonid immune repertoires (Guslund et al., 2020; Perdiguero et al., 2024). We also identified evidence of of phagocytic B cells (Wu et al., 2020): Cluster NA2 concomitantly expresses B-cell markers and Neutrophil markers, clusters with B-cells, and may represent this cell type, although we cannot rule out the role of doublets (two cells within the same droplet) in producing this signal given that B-cells physically interact with phagocytes during antigen presentation.

Our data reveals a range of novel cell type markers and marker combinations for distinct sub-lineages, which will facilitate future studies. Due to extensive evolutionary divergence between salmonids and model organisms (from which most immune cell markers are derived), identifying cell subtypes in non-models generally and fish specifically is challenging. Only a few cell-type specific antibodies exist, and qPCR markers require extensive verification due to WGD and functional divergence. We successfully combined a range of methods: expression of prior markers and overlap of prior markers with cluster markers, GO-term enrichment, pseudotime analysis to infer terminally differentiated cells, and extensive literature searches. Several well-established markers turned out to be robust for broad cell lineages in trout, for example the CD79 genes are useful markers of the B-cell lineage. In addition, we found numerous novel markers, some with no reported immune function, that were able to distinguish both broad immune cell lineages and sub-lineages. These include high choriolytic enzyme (HCE), an enzyme otherwise involved in digestion of the chorion during hatching, marking non-terminally-differentiated neutrophils, or *epdl1* (ependymin) as a new marker of a specific macrophage subpopulation (M3), which was previously reported expressed from the teleost central nervous system (McDougall et al., 2018). These and other markers reported in the supplementary data will be of great value in research and diagnostic settings and could potentially be used to produce better and species-/salmonid-specific antibody reagents for FACS.

Our analysis provides strong leads for in-depth evolutionary studies of gene duplications and their functional consequences. Salmonid genomes have retained intricate patterns of ohnolog deletion, pseudogenization, maintenance, and divergence (Gillard et al., 2021; Dimos & Phelps, 2023; Richman et al., 2024; Taylor et al., 2025) after the SSR4 whole genome duplication event, which are understudied in relation to the immune system. The fate of SSR4-derived duplicates has been debated, with Sandve et al (2018) concluding that the prevailing fate of ohnologue pairs is differential purifying selection (Sandve et al., 2018). The copy with more relaxed selection, in turn, may undergo neofunctionalization or pseudogenisation (indeed, this has been posited as the terminal fate of duplicated genes (Rastogi & Liberles, 2005)). Asymmetric expression patterns could signify such a transition, and indeed, half of 176 ohnolog pairs expressed in immune cells showed divergent expression patterns. A ‘gain or loss’ pattern is the most common, but also mutually exclusive patterns were observed. Interestingly, several cases involve partition of expression between fundamentally distinct lineages (e.g. B-cells versus Neutrophils, or B-cells versus Natural Killer Cells). This is consistent with subfunctionalisation, but could also reflect a transition state and, in millions of years, lead to the emergence of new function or to pseudogenisation. We did not further investigate the underlying sequence of coding regions or promotors, and also cannot distinguish divergence of ohnologue function from differential isoform usage. Our data are however an excellent starting point for in-depth cross-salmonid follow-up studies.

Finally, we find that molecular phenotypes correspond to rearing environment, which is of both conservatory and economic relevance. Commercial fish rearing environments have profound effects on fish health and disease resistance, with little knowledge regarding the underlying molecular mechanisms (Behringer et al., 2020). Yet, millions of fish are annually farm-raised and released into the wild for stocking purposes and are expected to support declining wild populations. Our data aligns with previous observations on the lower fitness of fish from more artificial rearing environments (Araki et al., 2008; Saura et al, submitted), and reveals that immune cells can indeed be associated with the raising environment based on their expression profile. Terminally differentiated Neutrophils, the majority of Macrophages, and naïve B-cells all differ depending on whether they were raised for at least two generations in a farm (farm), for a single generation on a farm (mix), or exclusively in natural streams (wild). In particular, genes involved in G protein-coupled receptor activity – a function closely linked with immune modulation, for example interleukin production (Saroz et al., 2019) and TLR signaling (Sharma et al., 2013) – were differentially expressed in two of the four tested clusters. G-protein coupled receptors have also been suggested as “domestication genes” that were repeatedly under selection during the domestication of pets and farm animals (Kleinau et al., 2024). The recurrence of similar gene sets across multiple cell types could imply that fixed to-be-discovered polymorphisms at cis- or trans-regulatory elements may affect the expression of these genes in a systemic fashion depending on the rearing environment. In addition, our setup also revealed that an entire T-cell cluster (T7) was present exclusively in fish of the ‘wild’ group. Given that these individuals derived from rivers known previously to harbor fish pathogens (namely *Tetracapsuloides bryosalmonae*), this origin-specific cell cluster may reflect prior infection history (for example, they may represent memory T-cells) and could in the future serve as an indicator for previous exposures to this or other pathogens. In summary, these findings warrant continued research on the origin of the observed transcriptional differences: Do they arise from environmental influences during early growth, or are they a consequence of natural selection in a farm rearing environment?

A curious and yet not entirely explained aspect of our data is the differential regulation of hemoglobin gene homologues in the wild environment in non-red blood cell clusters. While surprising, the production of hemoglobin by nonerythroid cells is not unheard of (Ghosh et al., 2018) and there is now evidence for a role of heme in a variety of cellular processes, including potentially immune responses via its interaction with NO (Dunaway et al., 2024). Another plausible explanation is the contamination of target immune cell libraries by RNA from red blood cells in the processed material which, although enriched for immune cells nevertheless contained abundant RBCs. Indeed, the expression of hemoglobin homologues in neutrophils, macrophages and B-cells was highest in the ‘mix’ sample B2 (**Fig. 10**) which also had the highest proportion of RBCs (around 16%, see **Fig. S5**).

## Conclusions

Our study delivers the first single-cell atlas of the brown trout immune system, revealing a complex and conserved immune architecture, unexpected cell-type markers, and profound impacts of genome duplication on immune gene evolution. By linking long-term evolutionary processes with short-term anthropogenic pressures, we expose how artificial rearing environments can rewire immune transcriptional programs—raising critical concerns for fish conservation and reintroduction strategies. Hatchery rearing, even for a single generation, leaves a detectable molecular imprint on the immune systems of brown trout—an apex predator and flagship species in freshwater ecosystems. The altered gene expression profiles in key immune cell types raise serious questions about the fitness and resilience of stocked fish, which are routinely released into the wild to support declining populations. As climate change and environmental degradation continue to erode natural habitats, it is imperative that stocking strategies evolve beyond numbers alone. We advocate for integrative policies that consider immunological integrity and genetic diversity when designing hatchery protocols and reintroduction programs. Investing in immunogenomic monitoring and promoting practices that mimic natural conditions could significantly enhance the long-term success of conservation and fisheries efforts. In this sense, our findings not only deepen our understanding of immune diversification in a key vertebrate lineage but also provide a valuable framework for future studies on functional immunity, aquaculture practices, and vertebrate genome evolution.

## Materials and methods

### Study design, sampling

Brown trout from different rearing histories and environments were used: wild group (wild), mix group (mix) and farm group (farm). The wild group consisted of young-of-the-year (≤ 1 year old) brown trout (F1 generation) that were fished during spring 2021 in two Swiss rivers (Brübach, coordinates between 47.471195 °N/ 9.136465 °E and 47.473303 °N/ 9.140526 °E; and Rörlibadbach between 47.473019 °N/ 9.136563 °E and 47.478731 °N/ 9.134755 °E. The parents (F0 generation) were born and raised in the wild. These waterbodies are known to be positive for Proliferative Kidney Disease, caused by a parasite *Tetracapsuloides bryosalmonae*, for generations (own observations). Although all animals were tested negative for T. bryosalmonae according to Bettge et al (Bettge et al., 2009), we cannot exclude that the F0 generation has faced an early infection or contact with *T. bryosalmonae*. The mix group (mix) originated from wild parents (F0 generation), female and male adult brown trout, which were fished in the same river system (Brübach) and spawned in autumn 2020. Fertilized eggs were transferred to a farm facility where they (F1 generation) were raised until spring 2021. The farm group (farm) originated from a Swiss farm, where two generations of brown trout were raised under farm conditions. Both mix and farm F1 generations were raised in the same hatchery facility (*Fischereizentrum Steinach*, Switzerland) from the stage of fertilized eggs until approximately three months of age-old fry. All animals used had the same genetic origin.

All F0 generations were transferred to the Institute for Fish and Wildlife Health (FIWI), University of Bern, Switzerland in May 2021, when fish were approximately 3 months of age. Each group of fish (wild, mix, farm) was distributed into six aquaria with equal fish numbers (n = 27 per aquaria). Each group consisted of three biological replicates. At arrival, two fish per aquarium were subjected to a health check, no pathogens were detected.

Fish were maintained at constant 16 °C with a flow-through water system and 12:12h light:dark photoperiod. The fish were fed a commercial diet twice a day with 2.5% of their body weight and additionally with live artemia in the first month after arrival. The fish were acclimatized to laboratory conditions for four months prior to sampling. Experiment was approved by the cantonal veterinary office (Bern, Switzerland) (Authorization number BE25/2021).

### Cell enrichment, library preparation and sequencing

Immune cells were isolated from the brown trout head and trunk kidneys when they reached the age of approximately 8 months. Both head and trunk kidneys are centres of hematopoietic cell production, and therefore material from both organs were pooled for each biological replicate. Fish were euthanized with an immersion bath in overdosed MS222 (150µg/l tricaine methanesulfonate; MS 222®, Pharmaq). The kidneys were then dissected and processed for RNA sequencing as follows:

The kidney tissue was passed through a 105µm pore size mesh filter with Leibovitz’s (L-15) medium (Thermofischer, Reinach, Switzerland) containing 10% fetal calf serum (FCS). The resulting cell suspension was then layered onto a sterile, isotonic Ficoll gradient (Ficoll-paque Plus, GE Healthcare Bio-Sciences AB, Sweden (with a density of 1.077 g/ml and spun at 750 x g for 40 minutes at 4°C to dissociate red blood cells from white blood cells. Leukocytes at the Ficoll/ medium interphase were aspirated, washed in L-15 medium and centrifuged at 290 x g at 4°C for 10 minutes. The cell suspension was aspirated and resuspended in PBS and kept on ice.

Cell suspensions were checked for viability and concentration using an automated cell counter (DeNovix® CellDrop Automated Cell Counter, NC, USA). The samples were diluted to a concentration ranging between 700 and 1900 cells/µL, depending on the sample. The cell suspensions were sent to the Bern Genomics Facility for 10x Chromium sequencing library preparation. All samples were prepared to ensure a target cell recovery of 10,000 cells and were subjected to cell isolation on the 10x Genomics Chromium Controller instrument. To minimize cell death, all steps, including PBL isolation, sorting, and Chromium™ Single Cell isolation, were completed on the same morning.

A total of 2500 cells per sample were loaded into the chips of the Chromium^TM^ Single Cell 3’ Gel Beads Kit (Single Cell 3’ v3.1 Dual Indexing) (10x Genomics) and processed on the Chromium Controller instrument to generate singe cell Gel Bead-In Emulsions (GEMs) following the manufacturer’s instructions.

Next, GEMs were subjected to library construction using the Chromium^TM^ Single Cell 3’ Library Kit v3.1 (10x Genomics). In the first step, reverse transcription was performed, generating cDNA tagged with a cell-specific barcode and a unique molecular index (UMI) per transcript. Fragments were size-selected using SPRIselect magnetic beads (Beckman Coulter, Brea, CA, USA). Illumina sequencing adapters were then ligated to the size-selected fragments, and cleaned up using SPRIselect magnetic beads. The quality of the final library was assessed using an Agilent 2100 Bioanalyzer (Agilent technologies, Amstelveen, The Netherlands). The samples were subsequently sequenced using a NextSeq 550 instrument (Illumina, CA, US) with 150PE chemistry.

### Augmented gene annotation

To maximise the amount of usable scRNAseq reads, which depends on the proportion of reads that can be confidently aligned to the transcriptome, we augmented the existing fSalTru1.1.104 ENSEMBL annotation using two published bulk RNAseq datasets derived from *S. trutta* kidney tissue (Ahmad et al., 2021; Sudhagar et al., 2019). Transcripts were assembled using a combination of de novo and genome guided approaches and further processed using the Mikado pipeline (Venturini et al., 2018) to identify a nonredundant set of transcripts not represented in the prior annotation. This approach enabled us to add 15,095 additional transcripts and 1,200 additional gene models. For more details please see the supplementary methods. The additional transcript and gene models (‘Mikado-derived genes’) were appended to a modified version of the ENSEMBL gtf (fSalTru1.1.104) which had been filtered to retain only protein-coding genes, lncRNAs, and the IG- and TR- genes that undergo somatic recombination before transcription. The GTF file of additional (non-ENSEMBL) genes and transcripts along with the functional annotations of Mikado-derived genes can be found in **Supplementary File 2**.

### scRNAseq read alignment and quantification

Raw FASTQ files from each of the nine samples were processed using CellRanger version 6.0.1 (10x Genomics, Inc.). The genome (fSalTru1.1, GCF_901001165.1) was indexed using ‘cellranger mkref’ with the augmented annotation file (see above), and reads were aligned to the genome using ‘cellranger count’ using non-default parameters –nosecondary and --include-introns, and otherwise default parameters.

### Processing of count data and clustering

The filtered barcode matrices from CellRanger-count were processed with the Seurat R package (Butler et al., 2018), using the Reciprocal Principal Components Analysis (RPCA) approach to normalize expression estimates across the three sample collection batches. In the RPCA approach, normalization (by SCTransform) and PCA were initially performed separately on samples from each of three sampling batches before integrating according to anchor features. To avoid subsequent clustering being overly influenced by cell cycle stage, the anchor features upon which PCA was performed were filtered to remove genes with annotation related to ‘cell cycle’ (using information from ENSEMBL). Similarly, given that red blood cells likely contributed a significant source of variation, hemoglobin genes were also excluded from the PCA. However, genes excluded from PCA were retained in the final set of 2000 variable genes. We did not explicitly filter according to mitochondrial gene expression which is liable to have retained dying cells in the dataset (Ilicic et al., 2016), although we note that mitochondrial filters may also exclude viable cells (Yates et al., 2025), particularly given the role of mitochondrial oxidative burst in some immune responses (Morris et al., 2022). For clustering, the FindNeighbors() function was run using the first 30 principal components. We then tested a series of resolution options for the FindClusters() step, from 0.1 to 1 in intervals of 0.1. Using clusters from all resolutions we then obtained a clustering tree using the clustree() function from the clustree package (Zappia & Oshlack, 2018), and obtained the SC3 index, an indicator of cluster stability, for each resolution. A resolution of 0.3 was found to have the highest SC3 value (**Fig. S8**) and we therefore chose this resolution for initial clusters (**Fig. S1**), but later performed additional subclustering of specific clusters in cases where subpopulations could be clearly distinguished from the expression of lineage markers (**Fig. S9**). No clear batch effects were evident from UMAP projections (**Fig. S10**).

Differential expression between clusters was performed by running the FindAllMarkers() function using the raw (non-integrated) expression values (given that the integrated values are statistically interdependent), producing a list of expression markers for each cluster (hereafter ‘cluster markers’). For further analyses we principally considered cluster markers with log2FC >/= 1 and which were detected in at least 60% of cells of a given cluster (‘high abundance’ markers), except for when clusters could be notably distinguished by expression patterns restricted to a subset of cells in the cluster.

### Assigning putative cluster identities

For assignment of clusters to broad immune cell classes, we examined the cluster-specific expression of putative cell type marker genes, a list of which was compiled a-priori. These genes (hereafter ‘prior markers’) were obtained by compiling a list of putative cell lineage markers from other species (humans in addition to some fish-specific markers), derived from the literature (**Table S1**). Putative brown trout homologues of these prior markers were identified by queries to the ENSEMBL database using the biomaRt R package (Durinck et al., 2005) to obtain putative brown trout homologues in combination with reverse blast searches (ENSEMBL blast tool, TblastN, default parameters) towards the brown trout genome using human queries and manually annotated fish queries from the literature. Reverse BLAST hits were aligned towards human and mouse reference sequences and query sequences to evaluate the overall degree of amino acid conservation (MEGA7, clustalw aligner). A simple neighbor-joining tree (MEGA7, Poisson substitution model, pairwise deletion) was constructed for markers with more pronounced sequence divergence (data not shown). In cases where homology could not be determined the marker was not used. This yielded in a list of 283 homologues of 129 prior markers of major cell lineage groups (e.g. B-cells, neutrophils) and functional classes (e.g. antigen presenting cells, cytotoxic cells, proliferating cells). For prior marker homologues which were present amongst the 2000 variable genes, expression values were summarised across markers belonging to the same lineage group or functional class using the AddModuleScore() function in Seurat, deriving expression modules corresponding with each major lineage group or functional class. We note that the list of prior marker genes was intended as a starting point and was not intended to be an exhaustive list of putative markers.

After grouping the clusters into broad immune (or other) cell types (i.e. neutrophils, monocytes / macrophages, T-cells, B-cells, and others), we used the cluster markers to further identify distinguishing gene expression features and more specific putative function. For this we considered only cluster markers that were unique to the broad cell type under investigation (e.g. markers of both T-cell and B-cell clusters were not used, although markers of ambiguous cluster N2 were allowed to be considered as either neutrophil or B-cell markers). We also performed GO term enrichment analyses on the cluster markers (all markers with log2FC >/= 1 and 60% detection in the cluster, with no criteria for uniqueness to a broad cell type) using the enricher() function from the ClusterProfiler package and GO terms of both ENSEMBL and Mikado-derived genes.

### Orthology analyses

To identify ohnologues amongst the 2000 variably expressed genes, *S. trutta* gene IDs were queried against the ENSEMBL database for homologues in the northern pike (*E. Lucius*) using the biomaRt package (Durinck et al., 2005). For genes with pike homologues, we filtered these to retain only those with a 2:1 ratio (trout:pike) and for which each member of the pair resided on a different trout chromosome. This resulted in a list of 152 putative ohnolog pairs amongst the variably expressed genes. UMAP plots for each pair were then visually assessed and assigned to one of the following four categories (similar to (Berthelot et al., 2014), but visually): 1) same pattern same levels (identical); 2) same pattern different levels; 3) gain/loss (with one ortholog expressed across a wider area than the other); 4) swap or partial swap (where orthologs have, at least partly, mutually exclusive expression patterns). Importantly, these assignments should be considered suggestive rather than final, particularly because gain/loss patterns may also result from differences in sequencing depth or sampling imbalances.

### Pseudobulk analyses

Clusters to use for pseudobulk differential expression analyses were chosen based on the following criteria: that they were clearly distinguishable from other clusters (i.e., abundant unique high abundance markers and not overlapping other clusters on a UMAP plot), and that they contained a reasonable cell count in all individuals (minimum 45). This led to the selection of clusters N1, M1, T5, and B3. The choice of B3 was further motivated by its identification as the pro/pre-B-cell cluster and thus alterations in this cluster could potentially have knock-on effects on subsequently differentiated B-cells. After pooling non-normalised read counts from each cluster / individual combination, the resulting pseudobulk count matrices were processed for differential expression analyses using the edgeR package (Robinson et al., 2009). Genes were first filtered to retain only those with at least 10 counts in at least one of the three origin groups (wild, mix, farm) prior to TMM-normalisation. PCA was calculated using the plotMDS() function which uses the top 500 most variably expressed genes by default. Differential expression was calculated using the estimateDisp() and glmFit() functions with a model design matrix that included group as the main factor while also accounting for experimental batch [model.matrix(∼batch+group)]. Genes were considered differentially expressed between groups at FDR < 0.05. GO term enrichment analysis (using the domain ‘molecular function’) was carried out using the enricher() function from the ClusterProfiler package (Yu et al., 2012) and GO terms of both ENSEMBL and Mikado-derived genes. For heatmaps, log2CPM values of differentially expressed genes were Z-score normalised and plotted using the Heatmap() function from the ComplexHeatmap package (Gu et al., 2016).

### Pseudotime analyses

The integrated, normalized data (SCT assay data with 2000 features for 83,847 cells) was converted into a ‘cell_data_set’-object with monocle (monocle3_1.3.4). After preprocessing the data (preprocess_cds) using the first 50 dimensions of the PCA a UMAP dimensionality reduction was performed. Cells were clustered with a resolution of 0.00001 which resulted in 24 clusters. This resolution was chosen because the number of cell clusters is close to the original 28. After running the ‘learn_graph’ function, subsets of clusters and starting points for trajectories were interactively selected based on prior biological knowledge. Two separate pseudotime analyses were performed: one on neutrophil clusters using the cluster corresponding with putative myeloid progenitors (cluster MP in main analysis) as the starting point and another on B-cells using the cluster corresponding with pre-/pro-B-cells (cluster B3 in the main analysis) as the starting point.

## Data and code availability

All raw sequencing data will be deposited to the NCBI sequence read archive (SRA) upon publication. Code for reproducing the core analyses can be found in the following github repository: https://github.com/jamesord/browntrout_scseq.

## Author contributions

Conceptualization – IAK, HS; Data Curation – JO; Formal Analysis – JO, SO, AB, MHS; Funding Acquisition – IAK, HS; Investigation (Experimentation, Data acquisition) HSM; Methodology (design and development) – HSM, HS, JO; MHS; Project Administration – JO, IAK, HS; Resources (provision of samples or materials) HSM; Supervision – IAK, HS; Visualization – JO, IAK, ST; Writing Draft – JO; Writing Review and Editing – IAK, HS, HSM, SO, ST, MHS

## Acknowledgements

We thank the Fish Warden of the Canton of St Gallen and the staff and management of Fischereizentrum Steinach (Canton of St Gallen) for assistance with fish acquisition, Pamela Nicholson of the University of Bern Next Generation Sequencing Platform for advice regarding sample preparation and for performing 10x Chromium sequencing, and Dragan Stajic for insightful discussion. Calculations were performed on UBELIX, the high-performance computing cluster of the University of Bern. OpenAI’s ChatGPT was used to assist in the completion of the manuscript, specifically to streamline some sections of the text (after initial drafting by the authors) and to check the consistency and completeness of the reference list.

