## Supplementary_material_v2 for "Single-cell analysis of a salmonid immune system (brown trout *Salmo trutta*) reveals evolutionary divergence and hatchery-induced transcriptional reprogramming": FiguresS1to10_Ord2025Trout.pdf

**Supplementary figure legends**

**Figure S1. (A)** UMAP projection of 83,847 cells grouped into 29 initial Seurat clusters using a clustering resolution of 0.3. **(B)** TSNE projection of the same clusters.

**Figure S2.** Gene expression indicator metrics of dead or dying cells for all 34 final clusters with boxplots superimposed on top of clouds of individual cells. **(A)** Module expression score (summarised expression metric) of six mitochondrial genes. **(B)** Number of expressed genes per cell, i.e. those with non-zero counts. **(C)** Log2 of the total read counts of all expressed genes per cell.

**Figure S3.** UMAP projections of Seurat clusters showing normalised expression values of genes that were found to be markers of multiple cell types, namely: **(A)** lyz had prominent expression in both neutrophils and macrophages, **(B)** CD74-like had prominent expression in both macrophages and B-cells, and **(C & D)** rpl12 and upf3b showed prominent expression in multiple lineages.

**Figure S4.** UMAP projections of Monocle clusters with cells coloured according to pseudotime value estimated using Monocle. Clusters are labelled according to the putative identity assigned to the Seurat clusters. **(A)** Pseudotime estimates of neutrophils and adjacent cell clusters, where the putative myeloid progenitor cluster (MP) was assigned as the starting point. **(B)** Pseudotime estimates of B-cells, where the putative pre-/pro-B-cell cluster (B3) was assigned as the starting point.

**Figure S5.** Plots for individual clusters of the fraction of cells assigned to the cluster in each individual sample of the wild, mix, and farm groups. Point shape corresponds to the experimental batch.

**Figure S6.** UMAP projections of Seurat clusters showing normalised expression values of two homologues of TRGC1-like, a marker of gamma-delta T-cells.

**Figure 7.** Pseudobulk differential gene expression profiles between fish of different origins, shown for cluster T5 (T-cells 5). **(A)**: Location of cluster T5 on the UMAP projection (left) alongside PCA plot derived from the top 500 most variably expressed genes (between individual pseudobulk samples) (right). On the PCA plot, point colour denotes individual origin (wild, farm, or mix), while point shape denotes the sampling batch. **(B)**: Heatmap of genes found to be differentially expressed in either the ‘farm’ or ‘mix’ group compared to the ‘wild’ group. Dendrograms represent hierarchical clustering of genes and samples according to the Z-scores of the Log2(CPM) values (cell colour intensity). Sample IDs (top of heatmap) are coloured according to the group from which the individual originated (‘Origin’). Gene names are given where available from the ENSEMBL database, otherwise the ENSEMBL gene ID is shown.

**Figure S8.** Mean SC3 stability index of Seurat clusters calculated for eleven values of *k* (clustering resolution). Higher values indicate that a cluster is more stable across multiple resolutions.

**Figure S9.** Rationale for finer-scale subclustering of three of the 29 initial Seurat clusters according to expression of canonical markers: **(A & B)** splitting of cluster 8 according to expression of CD8A (T-cell marker), **(C & D)** splitting of cluster 10 according to MARCO (Macrophage marker), and **(E & F)** splitting of cluster 19 according to the expression of the macrophage marker module (seven genes) which showed no expression in part of the cluster.

**Figure S10. (A)** PCA, **(B)** UMAP and **(C)** TSNE projections of 83,847 cells coloured according to experimental batch.

Figure S1

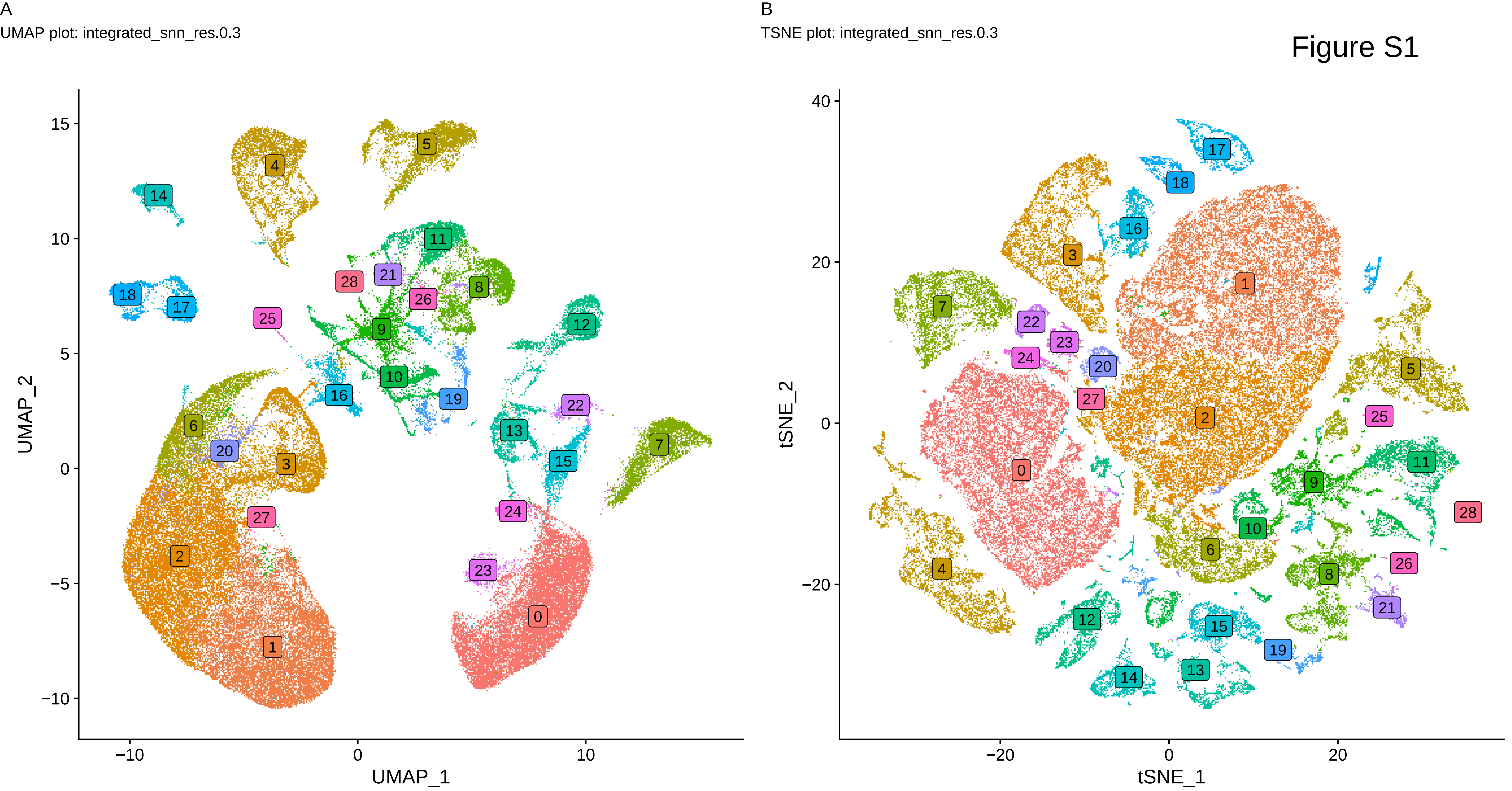
A
UMAP plot: integrated_snn_res.0.3
UMAP_2
UMAP_1
B
TSNE plot: integrated_snn_res.0.3
tSNE_2
tSNE_1

Figure S2

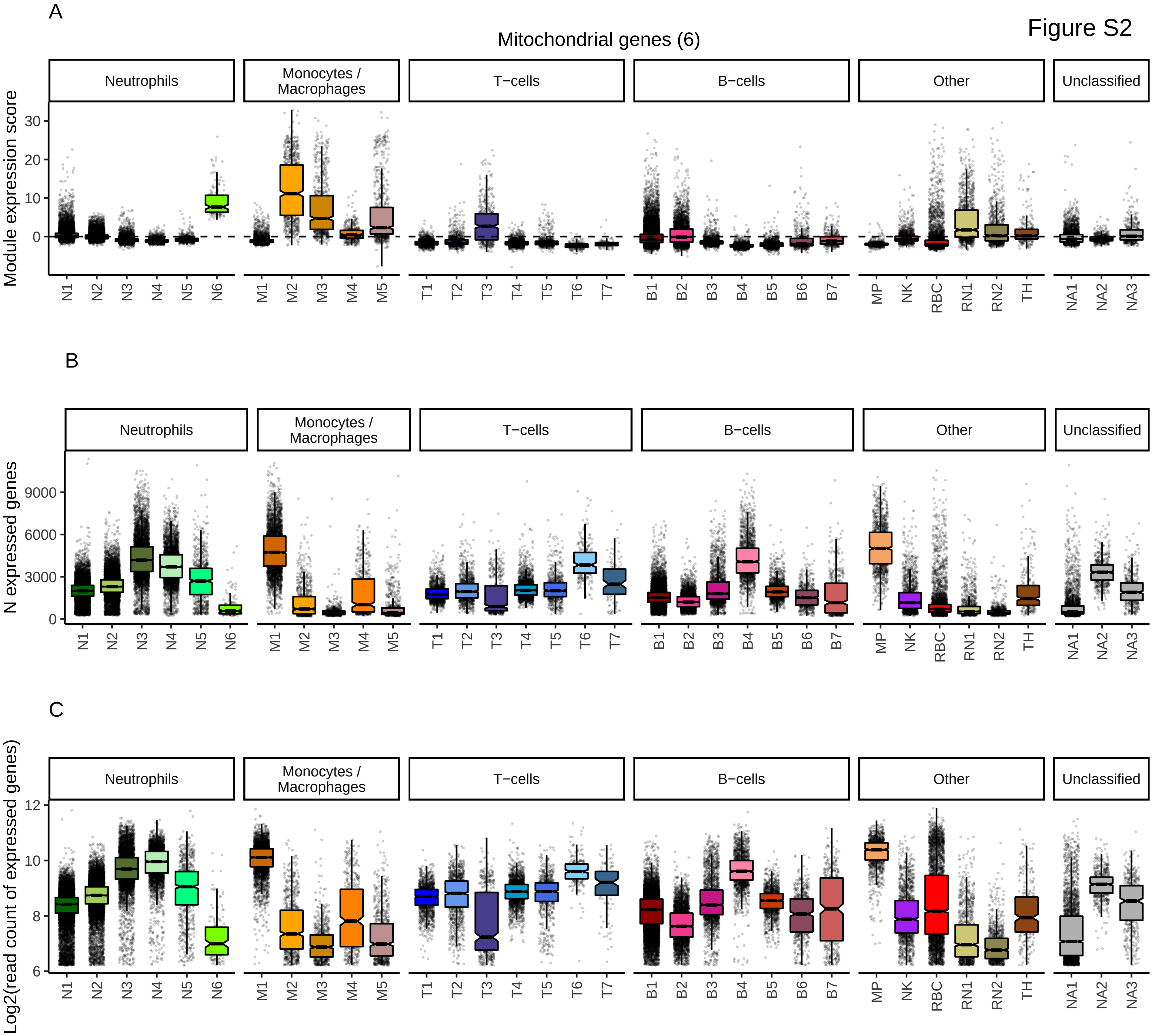
A
Mitochondrial genes (6)
Neutrophils
Monocytes / Macrophages
T−cells
B−cells
Other
Unclassified
Module expression score
N1 N2 N3 N4 N5 N6 M1 M2 M3 M4 M5 T1 T2 T3 T4 T5 T6 T7 B1 B2 B3 B4 B5 B6 B7 MP NK RBC RN1 RN2 TH NA1 NA2 NA3
B
N expressed genes
C
Log2(read count of expressed genes)

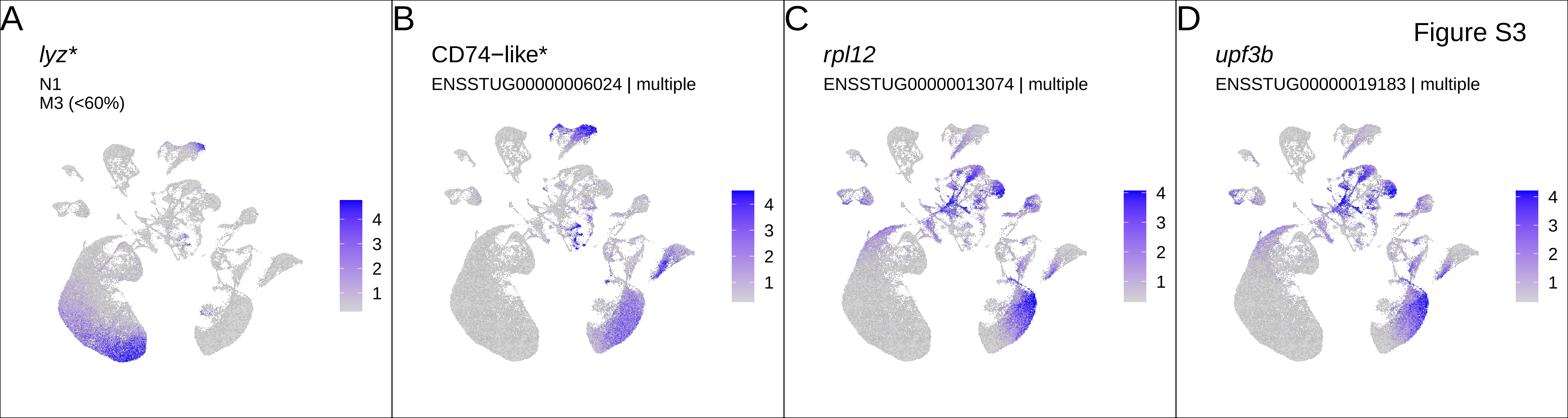

A
lyz*
N1
M3 (<60%)
4
3
2
1
B
CD74−like*
ENSSTUG00000006024 | multiple
4
3
2
1
C
rpl12
ENSSTUG00000013074 | multiple
4
3
2
1
D
Figure S3
upf3b
ENSSTUG00000019183 | multiple
4
3
2
1

Figure S4

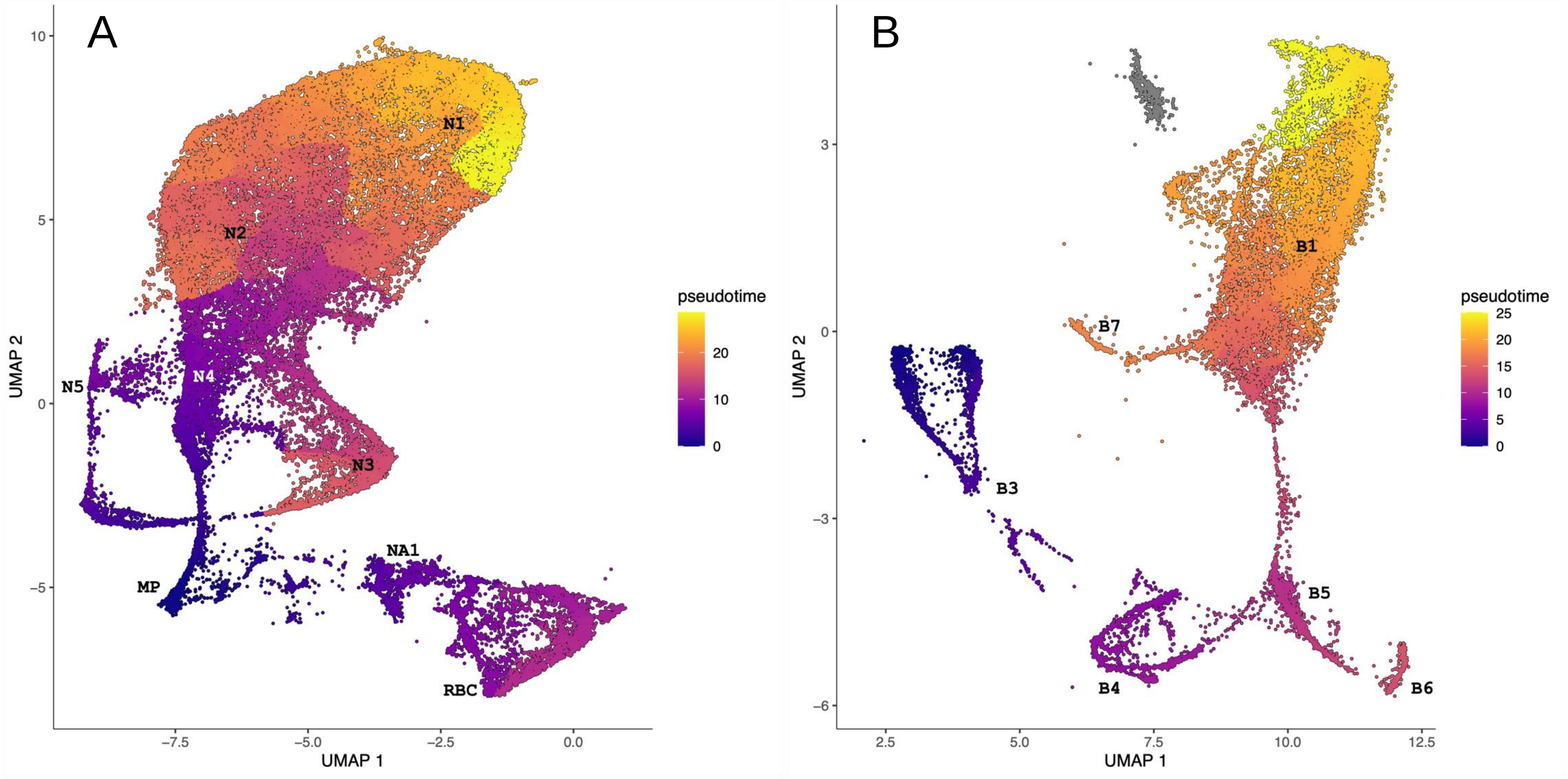

A
N1
N2
N3
N4
N5
MP
NA1
RBC
UMAP 1
UMAP 2
pseudotime
0
10
20
−7.5
−5.0
−2.5
0.0
−5
0
5
10
B
B1
B3
B4
B5
B6
B7
UMAP 1
UMAP 2
pseudotime
0
5
10
15
20
25
2.5
5.0
7.5
10.0
12.5
−6
−3
0
3

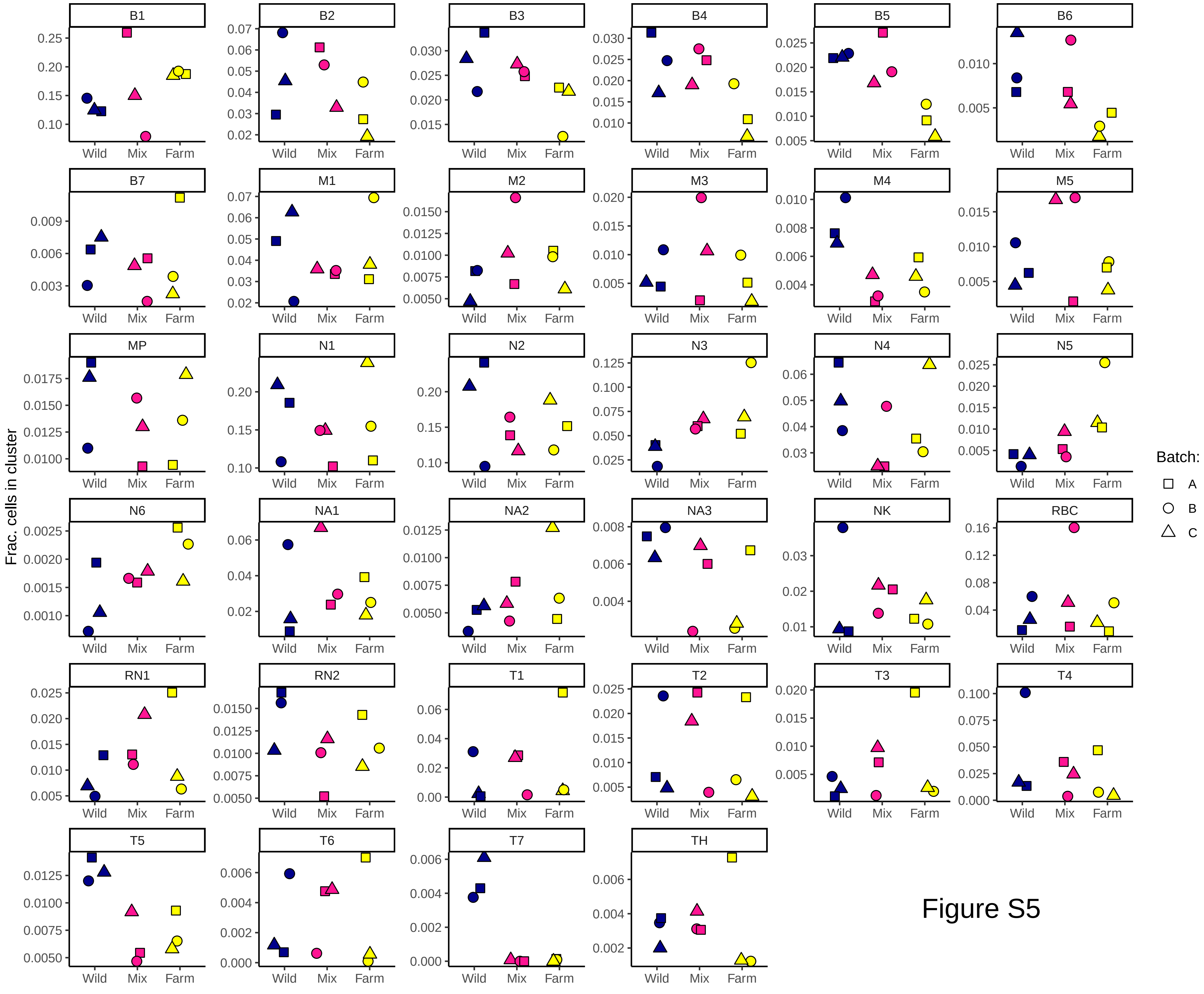

B1
B2
B3
B4
B5
B6
B7
M1
M2
M3
M4
M5
MP
N1
N2
N3
N4
N5
N6
NA1
NA2
NA3
NK
RBC
RN1
RN2
T1
T2
T3
T4
T5
T6
T7
TH
Wild
Mix
Farm
Frac. cells in cluster
Batch:
A
B
C


Figure S5

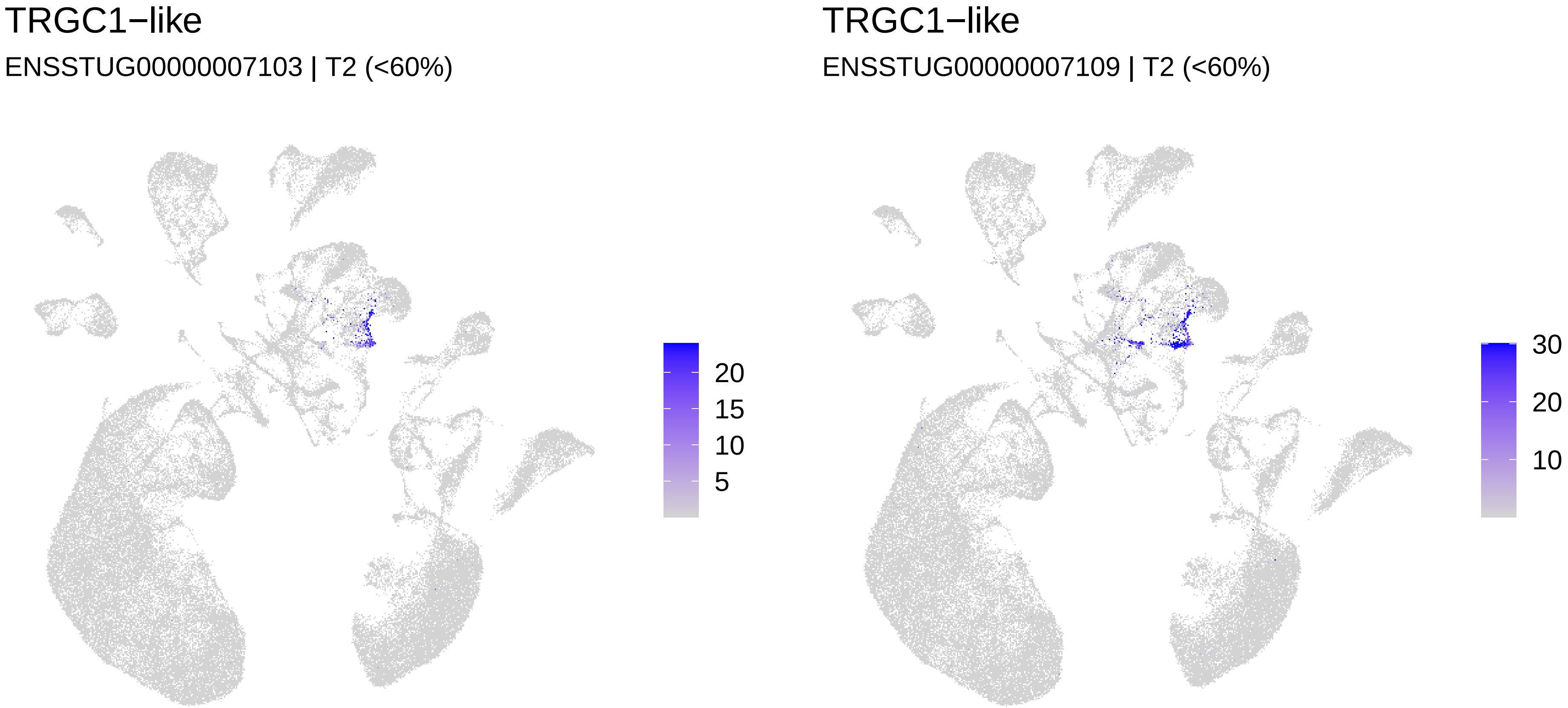
TRGC1−like
ENSSTUG00000007103 | T2 (<60%)
20
15
10
5
TRGC1−like
ENSSTUG00000007109 | T2 (<60%)
30
20
10

Figure S6

Figure S7

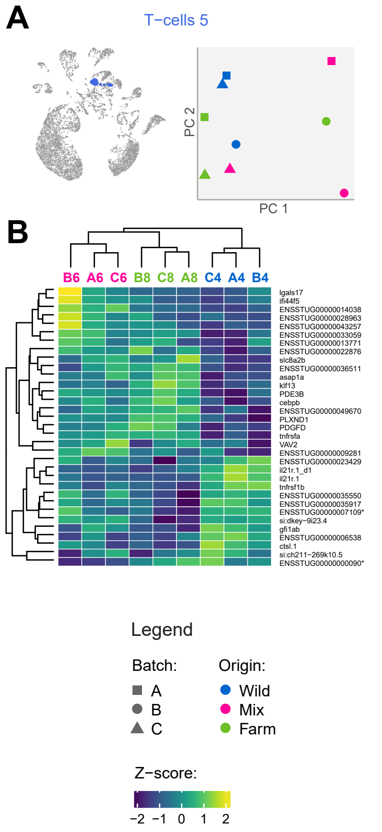
A
T−cells 5
PC 2
PC 1
B
B6
A6
C6
B8
C8
A8
C4
A4
B4
lgals17
ifi44f5
ENSSTUG00000014038
ENSSTUG00000028963
ENSSTUG00000043257
ENSSTUG00000033059
ENSSTUG00000013771
ENSSTUG00000022876
slc8a2b
ENSSTUG00000036511
asap1a
klf13
PDE3B
cebpb
ENSSTUG00000049670
PLXND1
PDGFD
tnfrsfa
VAV2
ENSSTUG00000009281
ENSSTUG00000023429
il21r.1_d1
il21r.1
tnfrsf1b
ENSSTUG00000035550
ENSSTUG00000035917
ENSSTUG00000007109*
si:dkey−9i23.4
gfi1ab
ENSSTUG00000006538
ctsl.1
si:ch211−269k10.5
ENSSTUG00000000090*
Legend
Batch:
A
B
C
Origin:
Wild
Mix
Farm
Z−score:
−2
−1
0
1
2

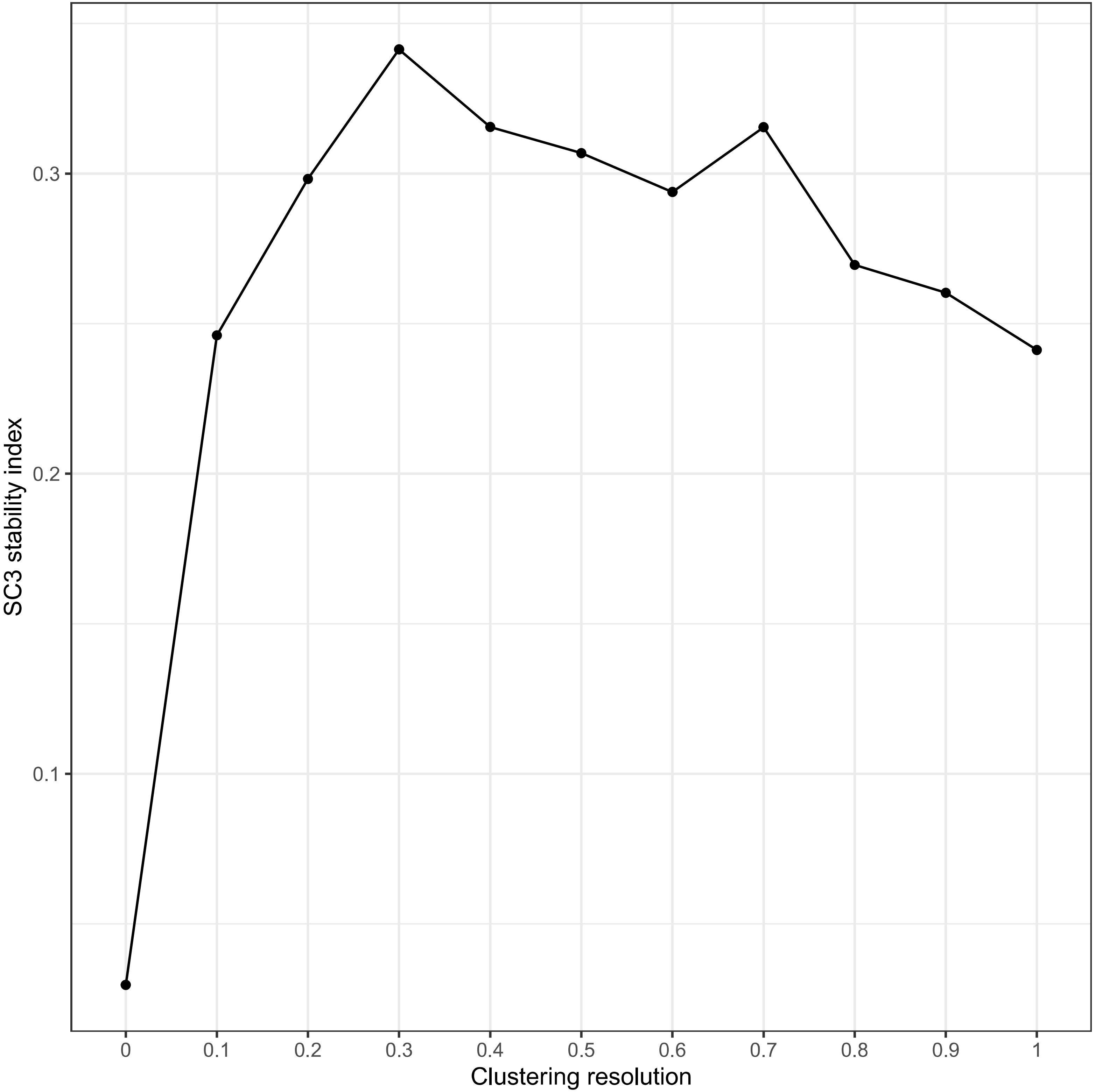

SC3 stability index
0.3
0.2
0.1
0
0.1
0.2
0.3
0.4
0.5
0.6
0.7
0.8
0.9
1
Clustering resolution

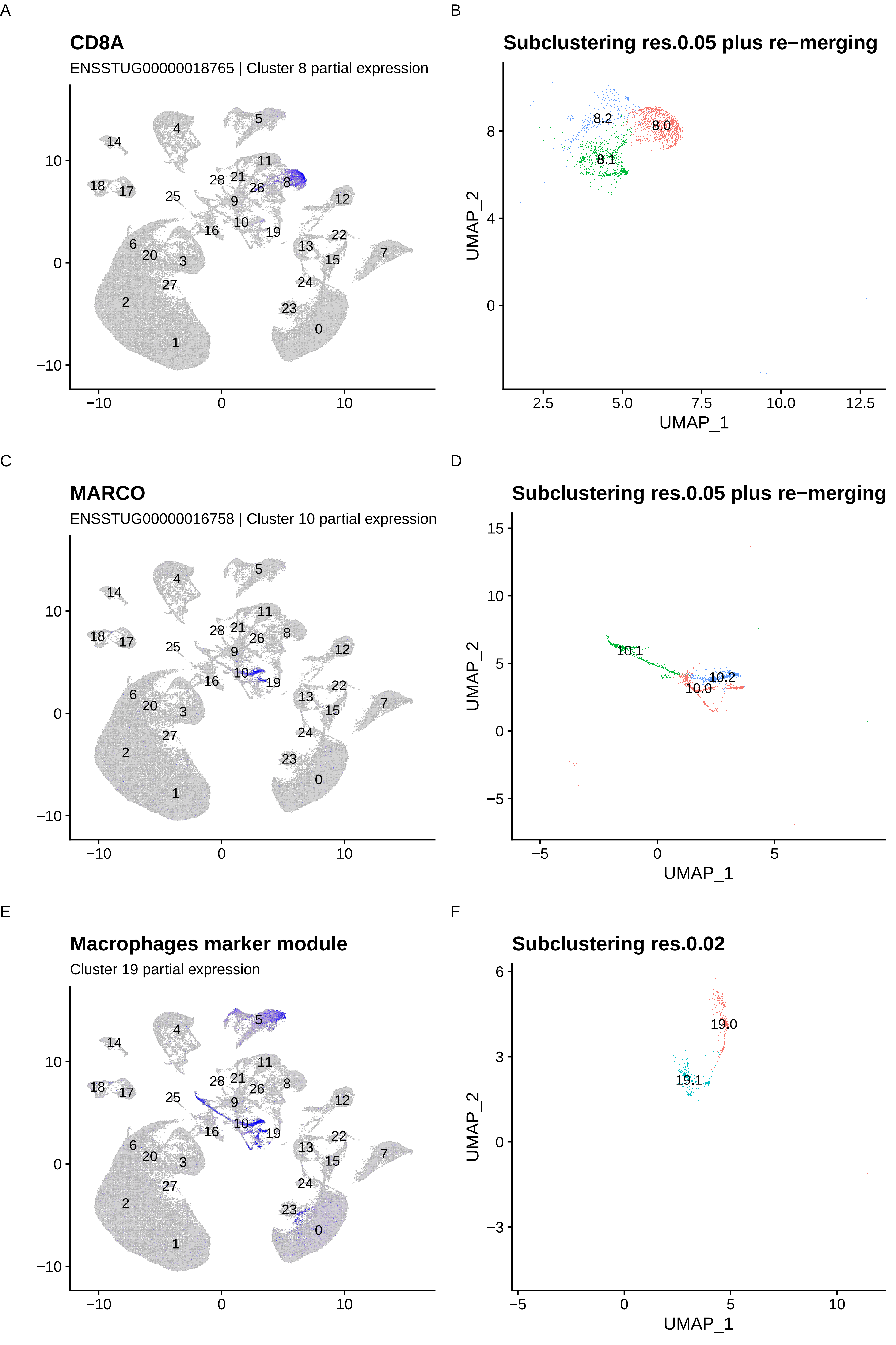
A
CD8A
ENSSTUG00000018765 | Cluster 8 partial expression
B
Subclustering res.0.05 plus re−merging
8.2
8.0
8.1
UMAP_2
UMAP_1
C
MARCO
ENSSTUG00000016758 | Cluster 10 partial expression
D
Subclustering res.0.05 plus re−merging
10.1
10.2
10.0
UMAP_2
UMAP_1
E
Macrophages marker module
Cluster 19 partial expression
F
Subclustering res.0.02
19.0
19.1
UMAP_2
UMAP_1

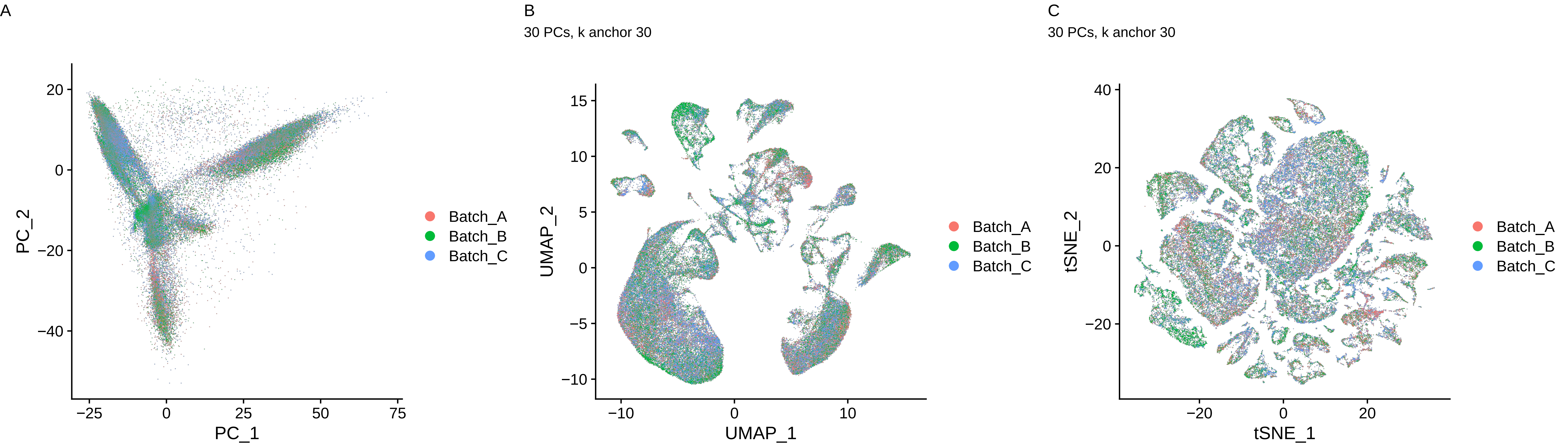
A
PC_2
PC_1
20
0
−20
−40
−25
0
25
50
75
Batch_A
Batch_B
Batch_C
B
30 PCs, k anchor 30
UMAP_2
UMAP_1
15
10
5
0
−5
−10
−10
0
10
Batch_A
Batch_B
Batch_C
C
30 PCs, k anchor 30
tSNE_2
tSNE_1
40
20
0
−20
−20
0
20
Batch_A
Batch_B
Batch_C
