## Supplementary_material_v2 for "Single-cell analysis of a salmonid immune system (brown trout *Salmo trutta*) reveals evolutionary divergence and hatchery-induced transcriptional reprogramming": SupMethods_Ord2025Trout.pdf

**Supplementary methods: Augmented genome annotation using bulk RNAseq data**

Although the current ESNEMBL transcriptome is of high quality (>98% complete BUSCOs), we opted to augment the existing annotations with a view to improving transcriptome mapping rates of 10x genomics reads (on average 44.8% of reads aligned confidently to the transcriptome on initial runs). Because read data generated from the 10x genomics 3' RNA assay are biased towards the 3' end of transcripts and therefore not ideally suited for full length transcript recovery, we used two published bulk RNAseq datasets derived from brown trout kidneys from: (1) twelve fish sampled from Austria (BioProject accession PRJNA542491 (Sudhagar et al., 2019)), totalling 417,050,173 100nt single-end reads, and (2) 14 fish sampled from Estonia (BioProject accession PRJNA668017 (Ahmad et al., 2021)), totalling 1,003,777,414 100nt paired-end reads.

Reads from each sample were trimmed using bbduk.sh from the BBMap suite v38.91 with parameters < ref=adapters ktrim=r k=25 mink=11 tpe=t tbo=t hdist=1 trimq=10 qtrim=rl >. Trimmed reads were filtered for rRNA using mirabait (MIRA v4.9.6) with default parameters, followed by alignment to the *S. trutta* genome fSalTru1.1 using Hisat2 v2.2.1. Splice site and exon files for use in index building and mapping were generated from the ENSEMBL gtf (annotation version fSalTru1.1.104) using scripts provided in the Hisat2 suite. Aligned reads from all samples were pooled and assembled using two de novo transcriptome assemblers, chosen for their ability to utilise existing reference information: Trinity v2.14.0 and RNAbloom v2.0.0. In its genome-guided mode, Trinity takes as input BAM files of reads aligned to a reference genome and restricts de novo assembly to reads that align to the genome in close proximity to one another. Trinity was run in genome-guided mode with the –max-intron-size parameter was set to 10000 and otherwise default parameters. Meanwhile RNAbloom was run using genome-aligned reads as input. While not using a reference genome, RNAbloom can take reference transcripts in FASTA format with which to augment the de Bruijn graph and guide transcript assembly, for which we used the *S. trutta* ENSEMBL cDNA FASTA file (from annotation version fSalTru1.1.104). Each of the two assemblers were run for each of the two abovementioned datasets separately, resulting in four de-novo transcriptome assemblies. To combine the de-novo assemblies, the FASTA files were concatenated and redundancy reduction was performed using the tr2aacds script from EvidentialGene v2022.01.20 (Gilbert, 2019) with parameter < -pHeterozygosity 7 > to account for multiple genotypes. This resulted in a set of 410,428 'okay' mRNA transcripts assigned to 140,155 clusters. Additionally, Stringtie v2.2.1 was run on each BAM file separately with default settings. Individual Stringtie-derived GTF files were subsequently merged by re-running Stringtie with the –merge option.

A set of consensus transcripts was derived by running the Mikado pipeline (Venturini et al., 2018) which used the aforementioned Stringtie GTF file, de-novo transcripts aligned to the genome with GMAP, and splice site predictions obtained using Portcullis. From the Mikado results, we identified additional transcripts of existing ENSEMBL genes as well as previously unannotated gene models, which were subsequently appended to the existing ENSEMBL annotation. In total we added 13,712 transcripts to

8,688 existing ENSEMBL genes and 1,383 transcripts of 1,200 ‘novel’ gene models. These novel genes were assigned putative functional annotation by extracting predicted protein sequences with TransDecoder v5.3.0 and subsequent BLASTP queries (assigning the closest BLAST hit as gene name). Predicted protein sequence queries were also submitted to the PANNZER2 web interface (Törönen et al., 2018) which assigns additional information including gene descriptions and GO terms. Annotations of novel transcripts in gtf format are provided in the supplementary .zip archive file.

Using our augmented annotations we were able to increase the average precentage of 10x reads aligned confidently to the transcriptome from 44.8% to 49.3%, further increasing to 57% when allowing intronic alignments.
